# Protify: A Benchmarking Platform Revealing Task Dependence and Homology Leakage in Protein Language Model Evaluation

**DOI:** 10.64898/2026.11.04.764373

**Authors:** Nikolaos Rafailidis, Logan Hallee, Tamar Peleg, Colin Horger, Luck Haviland, Jason P. Gleghorn

**Affiliations:** Center for Bioinformatics and Computational Biology, University of Delaware, Newark, DE, USA; Synthyra, PBLLC, USA; Department of Medical and Molecular Sciences, University of Delaware, Newark, DE, USA; Department of Biomedical Engineering, University of Delaware, Newark, DE, USA

**Author notes:** These authors contributed equally to this work (Nikolaos Rafailidis, Logan Hallee).

**Keywords:** Protein language models, Benchmarking, Protein property prediction, Zero-shot variant effect prediction, Homologyaware data splitting, Enzyme engineering, Open-source software

## Abstract

Protein Language Models (pLMs) have become essential tools for computational protein analysis, and the rapid proliferation of architectures, training objectives, and pretraining corpora has produced a diverse and expanding model landscape. Evaluation practices, however, remain fragmented: studies use different datasets, preprocessing pipelines, and reporting conventions, so results rarely transfer between publications. Here we introduce Protify, an open-source platform that standardizes pLM benchmarking across zero-shot, supervised, and transfer-learning settings through a unified, low-code interface. Protify currently supports 35 pretrained models and 61 curated datasets and it is easy to add custom versions of each. Leveraging Protify’s high throughput pipelines, we benchmarked 23 models on ProteinGym, discovering that DPLM3B is currently the top-performing self-supervised encoder. Another large-scale supervised evaluation across 32 diverse tasks, spanning enzyme classification, gene ontology annotation, protein-protein interaction, subcellular localization, and other functional properties, found that no single pLM dominated: 14 different models took the top score on at least one dataset, and the margins between the best and second-best models fell within run-to-run variability. On enzyme optimal pH prediction, a general-purpose probe pipeline matched or exceeded EpHod, a specialized model selected from 11,550 training runs, after 50 hyperparameter trials (*R*^2^ = 0.482 versus 0.425). Under a homology-controlled split of the same data, every pooled-embedding probe collapsed to near-zero *R*^2^, and only a transformer probe over full residue embeddings recovered appreciable signal (*R*^2^ = 0.248). Model rankings therefore depend as much on evaluation design as on the models themselves. Protify makes those design choices explicit, controlled, and replayable, and is freely available at https://github.com/Gleghorn-Lab/Protify.

## Introduction

Protein language models (pLMs), transformers trained on large corpora of protein sequences, have become standard tools for computational protein analysis. Representations learned from sequence alone predict structural and physic-ochemical properties (1–8), guide protein design (9–12), and enable functional annotation of uncharacterized sequences (13–17). The utility of pLMs has grown with machine learning progress and with the data behind them. Next-generation sequencing has driven exponential growth in protein sequence databases (18), with UniProt and MGnify now containing hundreds of millions to billions of sequences for diverse self-supervised pretraining (19, 20). In parallel, high-throughput methods like multiplexed assays of variant effect (21) and cell-free screening (22) supply an increasing number of labels for supervised learning. The resulting models differ in input data and training objective, spanning sequence-only transformers that learn from primary amino acid (3, 13, 18, 23) or codon (24, 25) sequences, models that incorporate evolutionary information (26–29), 3D structure (30, 31), or functional annotations (14, 32). Their objectives differ in what they encode: autoregressive training captures sequential dependencies, masked-language modeling uses bidirectional context, and contrastive objectives align representations with function or structure. Each choice shapes the learned representation, so performance varies across downstream tasks and datasets. Consequently, we suggest there is no single “best” pLM; model selection is therefore an empirical question (33).

Evaluation practices, however, remain fragmented. Studies use different datasets, preprocessing pipelines, evaluation metrics, and reporting conventions, so direct performance comparisons across publications are unreliable. Benchmark suites such as TAPE, FLIP and FLIP2, PEER, and PROBE address part of this problem by providing reusable tasks and standardized splits for protein function, structure, interaction, and fitness (34–38). More recent platforms have broadened this scope: PLMFit supports embedding extraction and transfer learning on user-provided datasets, PFMBench evaluates a broad collection of models and tasks, and Venus-Factory combines biological data retrieval, benchmarking, and modular pLM fine-tuning through command-line and graphical interfaces (39–41). Each resource makes different choices about model and dataset coverage, adaptation strategy, split construction, and reporting conventions, so results from one framework do not transfer directly to another. None of these resources combines frozen-embedding probes and parameter-efficient fine-tuning with built-in negative controls and experiment replay in a single interface.

We built Protify to make controlled comparisons routine and reproducible, with consistent preprocessing, training, evaluation, and baseline controls. Four design goals guided its development: (i) *modularity*, via a plugin architecture that supports rapid integration of newly released models and datasets; (ii) *systematic benchmarking*, through automated all-vs-all evaluations with built-in baseline controls and publication-ready figures; (iii) *accessibility*, through a well-documented command-line interface and a graphical interface for users who do not write code; and (iv) *reproducibility*, through extensive experiment logging, replay, and standardized reporting.

With these features, a single command runs thousands of model-dataset experiments, lowering the cost of principled model selection and production model development for drug discovery, enzyme engineering, and synthetic biology. We demonstrate Protify in three complementary case studies: zero-shot variant effect prediction on ProteinGym (23 models, 217 assays), large-scale supervised evaluation of frozen embeddings across 32 protein annotation tasks, and enzyme optimal pH prediction, where a general-purpose probe pipeline matches or exceeds EpHod, a model designed specifically for this task, on standard regression metrics (42). Under a homology-controlled split of the same dataset, all pooled-embedding probes collapse to near-zero *R*^2^, which bounds how much of the reported performance in this literature reflects generalization rather than sequence similarity.

## Results

### Training objective outweighs parameter count in zeroshot variant effect prediction

ProteinGym is the largest collection of deep mutational scanning (DMS) assays for predicting the functional effects of protein mutations. Its substitution benchmark comprises 217 DMS assays spanning diverse functional properties, protein families, and taxa, and has become a standard benchmark for zero-shot variant effect prediction (43). Each assay contains experimentally measured fitness values for amino acid substitutions at one or more sites, enabling direct comparison between model scores and experimental readouts. All models were scored using masked-marginal scoring with all mutated sites masked simultaneously, following Meier et al. (44) (see **Methods**).

Using Protify, we benchmarked a broad suite of bidirectional protein encoders, many of which have not been previously established on ProteinGym (**Table 1**): ESM2 models from 8M to 3B parameters, ProtBert-BFD^420M^, and the more recently released AMPLIFY, ESMC, GLM2, and E1 models (3, 13, 29, 45–47). We also included the emerging class of diffusion-based encoders (DPLM, DPLM2, and DSM), trained with a diffusion-based MLM objective that has been shown to yield stronger generative capabilities via iterative denoising while preserving, and in some cases improving, strong bidirectional representations for downstream predictive tasks (48, 49). This shared training objective let us apply a unified masked-marginal scoring protocol to every model except E1 (see **Methods**).

**Table 1.** ProteinGym substitutions benchmark results. Spearman correlations are shown for overall performance (Average) and stratified by functional property. Models are sorted in descending order by Average. Bold indicates best performance per column. Random-Transformer is a negative control: a randomly initialized ESM2_35M_ scored under the same protocol.

| **Model Name** | **Spearman Correlation** |  |  |  |  |  |
|---|---|---|---|---|---|---|
|  | **Average** | **Activity** | **Binding** | **Expression** | **Organismal Fitness** | **Stability** |
| DPLM<sub>3B</sub> | **0.421** | 0.427 | 0.340 | 0.422 | **0.395** | 0.520 |
| E1<sub>600M</sub> | 0.416 | 0.414 | 0.319 | **0.441** | 0.364 | **0.542** |
| E1<sub>300M</sub> | 0.414 | 0.438 | 0.324 | 0.430 | 0.345 | 0.533 |
| ESM2<sub>650M</sub> | 0.412 | 0.430 | 0.331 | 0.415 | 0.366 | 0.519 |
| DPLM2<sub>3B</sub> | 0.410 | **0.440** | 0.321 | 0.407 | 0.363 | 0.521 |
| DPLM<sub>650M</sub> | 0.407 | 0.413 | **0.344** | 0.409 | 0.354 | 0.514 |
| ESM2<sub>3B</sub> | 0.404 | 0.423 | 0.311 | 0.403 | 0.377 | 0.507 |
| ESMC<sub>300M</sub> | 0.404 | 0.422 | 0.307 | 0.408 | 0.360 | 0.521 |
| ESMC<sub>600M</sub> | 0.403 | 0.423 | 0.289 | 0.421 | 0.361 | 0.520 |
| E1<sub>150M</sub> | 0.398 | 0.426 | 0.316 | 0.420 | 0.302 | 0.527 |
| ESM2<sub>150M</sub> | 0.386 | 0.399 | 0.317 | 0.402 | 0.304 | 0.509 |
| DPLM<sub>150M</sub> | 0.379 | 0.385 | 0.322 | 0.403 | 0.288 | 0.498 |
| DPLM2<sub>650M</sub> | 0.378 | 0.411 | 0.295 | 0.403 | 0.308 | 0.471 |
| DSM<sub>150M</sub> | 0.377 | 0.386 | 0.316 | 0.392 | 0.291 | 0.502 |
| AMPLIFY<sub>350M</sub> | 0.359 | 0.373 | 0.214 | 0.364 | 0.390 | 0.455 |
| AMPLIFY<sub>120M</sub> | 0.353 | 0.393 | 0.226 | 0.385 | 0.363 | 0.400 |
| ProtBert-BFD<sub>420M</sub> | 0.351 | 0.353 | 0.291 | 0.383 | 0.304 | 0.424 |
| DPLM2<sub>150M</sub> | 0.337 | 0.356 | 0.296 | 0.379 | 0.245 | 0.411 |
| ESM2<sub>35M</sub> | 0.316 | 0.315 | 0.282 | 0.344 | 0.216 | 0.424 |
| DSM<sub>650M</sub> | 0.246 | 0.244 | 0.164 | 0.289 | 0.209 | 0.324 |
| ESM2<sub>8M</sub> | 0.219 | 0.191 | 0.250 | 0.266 | 0.139 | 0.252 |
| GLM2<sub>150M</sub> | 0.197 | 0.220 | 0.193 | 0.240 | 0.186 | 0.147 |
| GLM2<sub>650M</sub> | 0.197 | 0.245 | 0.208 | 0.283 | 0.195 | 0.057 |
| Random-Transformer | −0.040 | −0.025 | −0.023 | −0.031 | −0.052 | −0.067 |

Across this panel, DPLM_3B_ achieved the highest average Spearman correlation (0.421), placing it among the top performing pLMs with only self-supervised denoising and single-sequence input during inference (**Table 1**). The E1 models followed, consistently surpassing their size-matched ESM2 counterparts, and the two ESMC models matched ESM2_3B_. Protify closely reproduced previously reported ESM2, ESMC, and E1 performance across model sizes to within 0.007 of ProteinGym leaderboard values; we attribute these minor deviations to our use of simultaneous masking rather than the iterative masking approach used in the original ProteinGym and E1 repositories (see **Methods**).

DPLM_650M_ scored just below ESM2_650M_ at a similar parameter count, even though DPLM_3B_ outperformed its ESM2 counterpart. Notably, DPLM2 did not surpass DPLM at any parameter scale on average; however, it did achieve the highest score on the activity portion of the benchmark (0.440). AMPLIFY models scored below their size-matched ESM2 baselines, with AMPLIFY_350M_ falling below ESM2_150M_ despite having more than twice the parameters, and ProtBert-BFD_420M_ performing comparably to AMPLIFY. The diffusion-based families diverged: DPLM ranked near the top, whereas DSM scaled inversely, with DSM_650M_ scoring well below DSM_150M_. GLM2 models performed below every other architecture tested at both parameter scales. Architectural and objective choices therefore matter independently of scale: the E1 family surpassed size-matched masked language models at every size tested, whereas diffusion-based training in DPLM surpassed ESM2 only at 3B parameters.

### No single pLM dominates across 32 supervised protein tasks

Beyond mutation effect prediction, researchers also rely on pLMs for a wide range of downstream structure- and function-related tasks. To assess model performance across these tasks, we used Protify to run a many-model vs. many-dataset benchmark on 32 supervised protein property prediction datasets, which we call the vector benchmark. These datasets span diverse areas of protein annotation, including enzyme properties (EC numbers, *k*_*cat*_), Gene Ontology annotation, protein-protein interaction (PPI), subcellular localization, taxonomy, expression prediction, and other fitness measures (**Supplementary Table S6**). The benchmark measures how well each pLM’s representations predict distinct labels under a standardized multi-layer perceptron (MLP) probing protocol.

For each model-dataset combination, we extracted embeddings from the frozen model and pooled them as described in Methods. We chose mean-variance pooling over residue positions on the basis of a preliminary evaluation across a 10-dataset subset of the vector benchmark, where it ranked best on average; that evaluation used ESMC_300M_ embeddings and a random forest rather than the MLP probe applied here (**Supplementary Table S1**). We then trained a three-layer MLP probe using the concatenated embeddings as features, with the same hyperparameters and early-stopping criteria for each combination. We repeated training three times with different random seeds, and the reported scores are the mean performance. This protocol fairly compares the innate correlation of each model’s representations with relevant downstream tasks by holding architecture and training details constant while accounting for seed-to-seed variability.

In addition to the amino acid-based pLMs, we evaluated CaLM_86M_, a codon language model pretrained on nucleotide sequences (25). For the *ecoli_expression* dataset, which consists of DNA coding sequences, we evaluated CaLM_86M_ directly. For all other datasets, we reverse-translated sequences to DNA using the most frequent human synonymous codon for each amino acid (see **Methods**).

Performance was highly task-dependent across the 32 datasets, with no single pLM consistently topping the leaderboard (**Fig. 2**). DPLM_3B_ was the most frequent top performer, with the best score on 6 of 32 datasets, followed by ANKH_400M_ with four. In total, 14 different models were the best performer on at least one task. Notably, the margins separating top-performing models were remarkably thin. Welch’s *t*-tests across three random seeds show that the difference between the best and second-best model was not sta-tistically significant on any dataset (*p >* 0.05; **Table S4**). Three seeds provide limited statistical power; nevertheless, these results suggest that fine-grained ranking differences should be interpreted cautiously (see **Supplementary Note 5** for details and limitations). **Supplementary figure S2** shows per-cell standard deviations across the three seeds.

Several task-specific patterns stood out. CaLM_86M_, the only codon language model evaluated, achieved the highest MCC (0.14) on soluprot, a protein sequence dataset for which it relies on deterministic reverse translation to DNA (see **Methods**), a naive strategy that discards the synonymous codon variation CaLM_86M_ was trained to exploit. More strikingly, on *ecoli_expression*, the only DNA-sequence dataset in the benchmark where CaLM_86M_ could leverage its native codon-level context, it performed worse than any other pLM (MCC 0.23), while AMPLIFY_120M_ achieved the highest score (0.47). These findings suggest that predicting expression from sequence alone can be difficult, and that codon-level signal is not sufficient on its own for additional performance gains. ProtBert-BFD_420M_ led the taxonomy classification tasks, achieving the top score on *taxonomy_different* and *taxonomy_species*.

The two multi-input datasets constructed using C3 data splits over clusters (50, 51), KSMoFinder-clustered (a kinase-substrate phosphorylation task) and the Bernett PPI dataset, yielded lower and more variable performance across models. This indicates that homology cluster-based C3 splits alongside other types of metadata substantially reduce data leakage and produce more rigorous evaluations, consistent with existing literature (8, 51–53). This difficulty is not specific to pLM probes: among 10 classical classifiers trained on the same ESMC_600M_ embeddings, most scored at or near zero MCC and the best reached 0.17 (**Supplementary Note 6**).

Interestingly, DSM_650M_ ranked fourth overall in average vector benchmark score (0.534), making it the best-performing model below 1 billion parameters. However, on ProteinGym it performed poorly, with an average Spearman correlation of 0.246, well below much smaller models such as ESM2_35M_ (0.316). Such disagreements between leader-boards highlight pLMs’ uneven performance across evaluations, where different training objectives can improve or degrade performance along different axes.

### A general-purpose probe matches a specialized enzyme pH predictor

The all-vs-all benchmark reveals which models encode the strongest representations for a given task, but does not test the ceiling of those representations. A probe with selfattention over full-sequence residue embeddings can substantially improve performance, sometimes surpassing models engineered for a single task. We tested this on enzyme optimal pH (pHopt) prediction, a property central to enzyme engineering for industrial and therapeutic applications. EpHod (42) is a dedicated pHopt model, selected from an evaluation of 11,550 trained model instances spanning diverse representations and architectures. It ensembles two ESM-1v-based models: a residual light attention network pretrained on 1.9 million environmental pH annotations and fine-tuned on pHopt using per-residue embeddings, and a support vector regression model using mean-pooled embeddings.

During dataset preparation, we identified 24 identical sequences appearing across splits under different UniProt accessions, 20 between training and test and four between validation and test. Several of these pairs also carried conflicting pHopt labels for the same sequence. We therefore constructed the `optimal_ph_rigor` split, with zero cluster or sequence overlap between any pair of splits (8,015/430/439 train/validation/test; see Methods).

In the all-vs-all benchmark, DPLM2_3B_ achieved the highest *R*^2^ (0.393) of any model on the standard pHopt split using only an MLP probe on frozen mean-variance embeddings, approaching EpHod Ensemble’s *R*^2^ of 0.425 after seconds of training with no task-specific engineering (**Table 2**). We selected DPLM2_3B_ and trained a transformer probe on its full residue embeddings, selecting hyperparameters via a Bayesian sweep over validation *R*^2^ (see Methods). On the standard test set (*n*=1,971), the transformer probe reached *R*^2^ = 0.482, surpassing EpHod Ensemble on all five standard regression metrics (**Table 2, Fig. 3A,B**). On EpHod’s sample-weighted metrics, which upweight the sparse acidic and alkaline bins, the ordering reverses and the EpHod Ensemble leads (**Supplementary Note 2**).

**Table 2.**
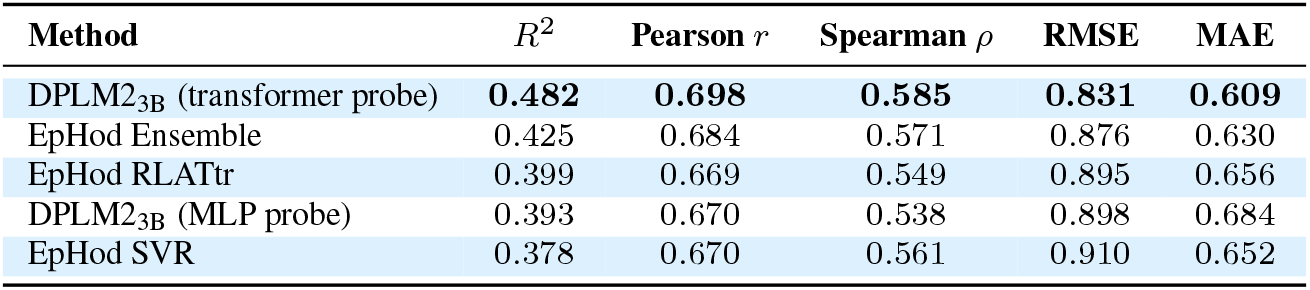
Comparison of regression metrics on the pHopt test set (*n*=1,971). DPLM2_3B_ results use the standard EpHod split. Bold indicates best performance per column. The MLP probe was trained on frozen DPLM2_3B_ embeddings from the all-vs-all benchmark; the transformer probe used optimized hyperparameters (see Methods).

| **Method** | *R*<sup>2</sup> | **Pearson** *r* | **Spearman** *ρ* | **RMSE** | **MAE** |
|---|---|---|---|---|---|
| DPLM2<sub>3B</sub> (transformer probe) | **0.482** | **0.698** | **0.585** | **0.831** | **0.609** |
| EpHod Ensemble | 0.425 | 0.684 | 0.571 | 0.876 | 0.630 |
| EpHod RLATtr | 0.399 | 0.669 | 0.549 | 0.895 | 0.656 |
| DPLM2<sub>3B</sub> (MLP probe) | 0.393 | 0.670 | 0.538 | 0.898 | 0.684 |
| EpHod SVR | 0.378 | 0.670 | 0.561 | 0.910 | 0.652 |

**Fig. 1.**
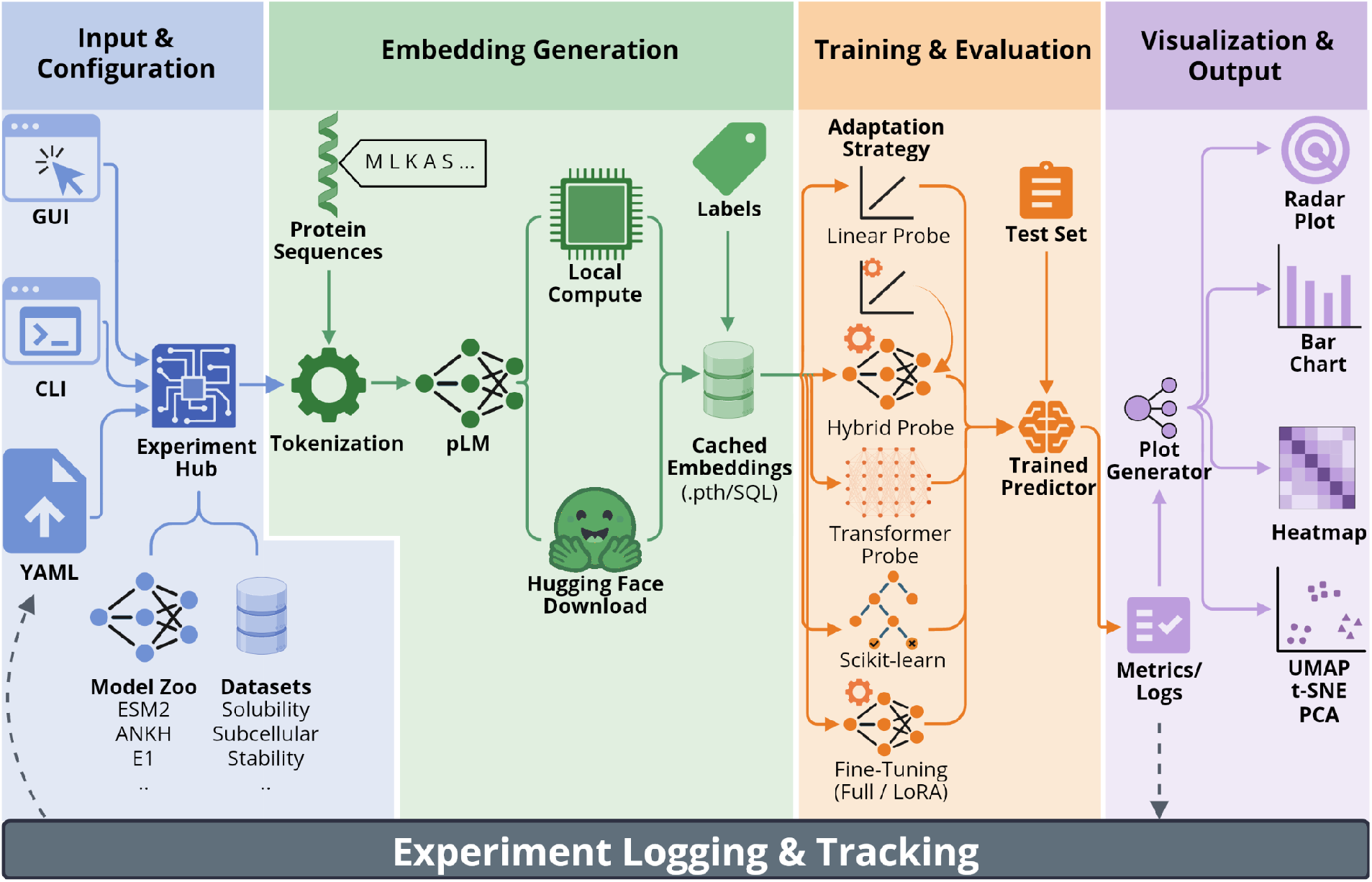
Protify workflow. **(1) Input & configuration:** users select models, datasets, and settings through a graphical interface (GUI), command-line interface (CLI), or YAML configuration file. **(2) Embedding generation:** Protify embeds each sequence with the selected pLMs, or downloads precomputed embeddings from Hugging Face, and caches them locally. **(3) Training & evaluation:** a downstream predictor is trained on the cached embeddings using the chosen adaptation strategies and evaluated on a held-out test set. **(4) Visualization & output:** Protify generates publication-ready figures and logs all metrics; passing the log file back to Protify replays the full experiment.

**Fig. 2.**
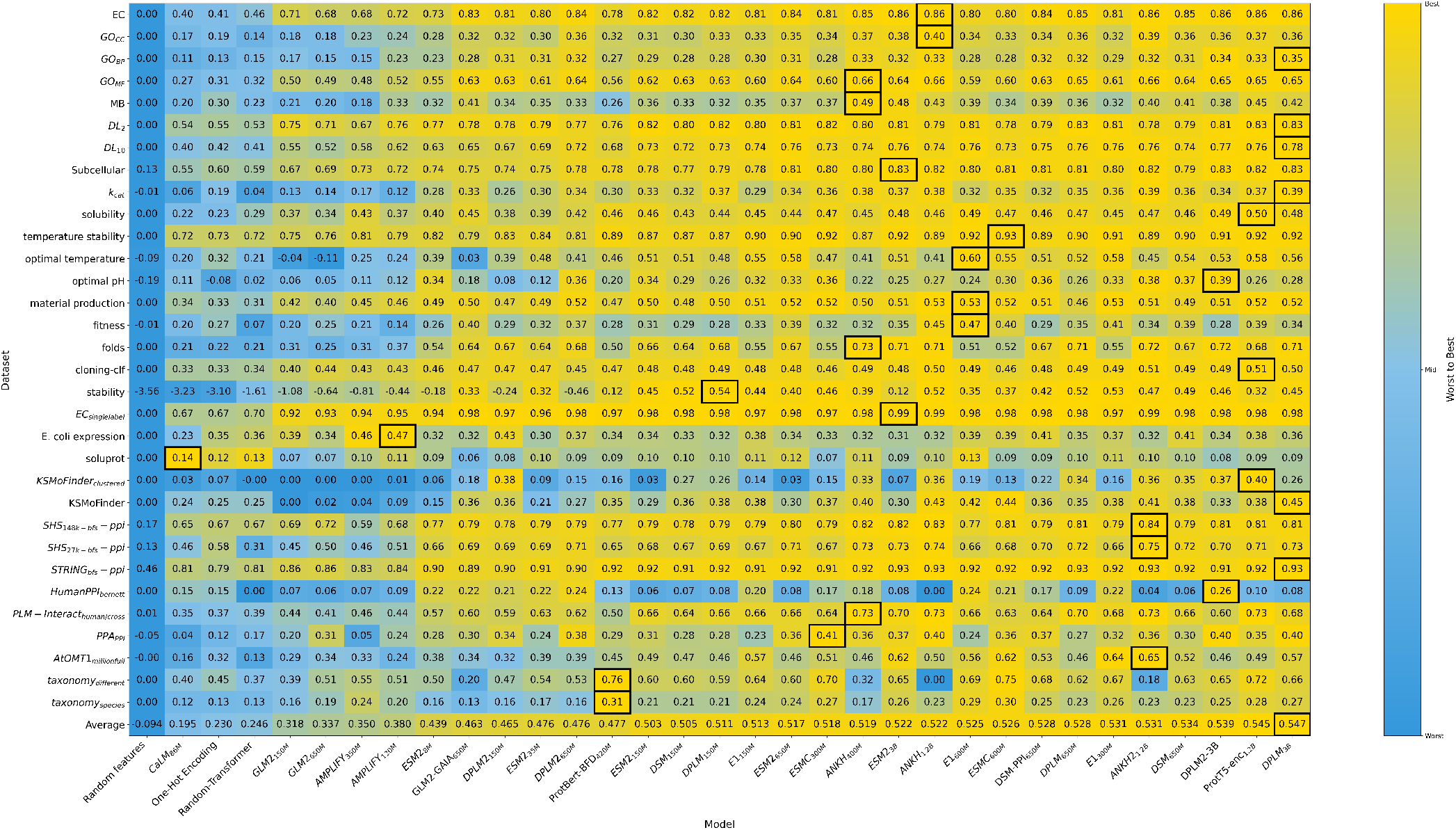
All-vs-all supervised vector benchmark across 32 protein annotation tasks. Heatmap of test set performance for each model-dataset pair under a fixed probe training protocol on frozen, mean-variance pooled embeddings. Each cell displays the average of the primary metric for that dataset across three random seeds (MCC for classification, *R*^2^ for regression). Colors are independently normalized per row, highlighting the worst (blue) to best (yellow) performing model within each task. The final row shows the average performance across all datasets. Models are sorted by average performance in ascending order from left to right.

**Fig. 3.**
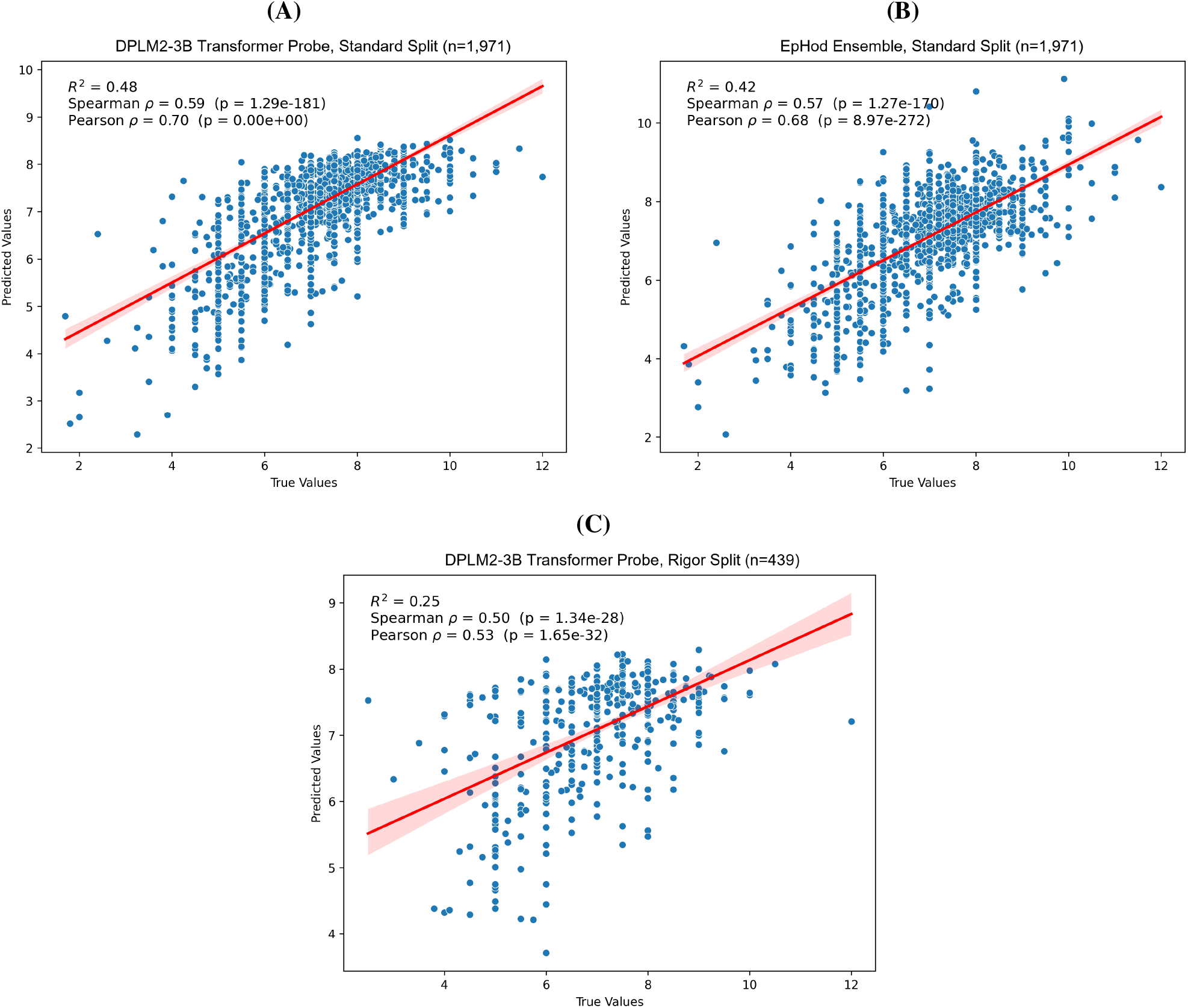
Enzyme optimum pH prediction. Scatter plots of predicted versus true pHopt values. **(A)** DPLM2_3B_ transformer probe on the standard test split (*n*=1,971). **(B)** EpHod Ensemble on the standard test split (*n*=1,971). **(C)** DPLM2_3B_ transformer probe on the homology-aware rigor split (*n*=439). Red line and shaded region in each panel show the linear regression fit and the 95% confidence interval. Metrics (*R*^2^, Spearman *ρ*, Pearson *r*) are annotated in each panel.

### Homology-controlled splits collapse frozen-embedding performance

On the rigor split, where partitioning by cluster at 30% sequence identity ensures no homologous sequences are shared across splits, all MLP probes failed, with *R*^2^ near or be-low zero and E1_300M_ scoring highest at *R*^2^ = 0.039. Frozen pooled embeddings alone therefore do not produce accurate pH predictions under strict generalization conditions. The DPLM2_3B_ transformer probe reached *R*^2^ = 0.248 on this split (**Fig. 3C**), whereas the MLP probe on the same backbone scored *R*^2^ = -0.075, near the random-transformer baseline. This gap suggests that per-residue positional information carries predictive signal for pH that pooling loses but full residue processing can recover.

## Discussion

Protein language models span a wide range of architectures, objectives, and pretraining corpora; our results show that these design choices do not produce a stable, universal ranking. Instead, the “best” model depends strongly on the task and evaluation protocol. Model selection therefore requires direct empirical comparison. Rankings from one evaluation framework (39–41) should not be assumed to hold in another.

PFMBench, for example, applies a 30% sequence identity threshold when constructing most supervised dataset splits, a choice that can substantially alter model performance and rankings. Protify does not remove this dependence on evaluation design; it holds those choices fixed while comparing candidate models side by side.

The underperformance of the AMPLIFY models on both benchmarks was unexpected: in the vector benchmark, they ranked near the bottom, above only models pretrained on different or mixed modalities. AMPLIFY was pretrained on UniRef100, which, unlike the commonly used UniRef50 and UniRef90 variants, retains near-complete sequence diversity on the premise that broader coverage of the natural sequence space may yield richer representations (45). Because ProteinGym assays consist of amino acid substitutions within known protein families, we expected AMPLIFY to perform competitively, having seen more examples of closely related sequences differing at individual positions. AMPLIFY_350M_ instead scored below the half-sized ESM2_150M_ on ProteinGym substitutions and on par with the much older ProtBert-BFD_420M_. Spinner et al., evaluating AMPLIFY models trained on successive UniRef100 releases, found that zero-shot ProteinGym performance does not improve monotonically with additional data and can degrade despite continued dataset growth (54). The authors partially attribute this to UniRef100’s increasing redundancy over time, as newly added sequences add duplicate existing ones rather than expanding true sequence diversity.

Across most model families in the vector benchmark, performance scaled monotonically with parameter count; ESM2, ESMC, GLM2, DSM, and ANKH all gained as model size increased. DPLM2 followed this trend less cleanly. Its 3B variant performed comparably to the size-matched DPLM_3B_ predecessor, but the 150M and 650M variants fell considerably further behind their DPLM counterparts, performing at the level of ESM2_35M_. We evaluated DPLM2 on sequences alone, without the structure tokens it was also trained on (55); even so, DPLM2_3B_ achieved the highest score on the challenging Bernett dataset. Among the E1 family, the 300M variant outperformed the 600M on average, despite the latter achieving the highest score on three tasks and the former on none. These observations suggest that, while parameter count is a useful first-order heuristic, it is not always a reliable proxy for downstream performance. The same holds on ProteinGym, where neither DPLM2_3B_ nor ESM2_3B_ beat ESM2_650M_, a model with roughly five-fold fewer parameters.

Both ProtBert-BFD_420M_’s strength on the taxonomy tasks and ProtT5_enc-1.2B_’s second-place overall finish, despite ProtT5_enc-1.2B_ being one of the oldest models evaluated, may trace to BFD pretraining: BFD comprises ∼2.1 billion predominantly metagenomic protein sequences, whose breadth could preserve coarse taxonomic cues that redundancy-filtered corpora discard, and ProtT5 pairs that coverage with non-redundant UR50 fine-tuning. This may partially explain this model’s durability. Many of the top models converge to a similar average performance (∼0.54), raising the question of whether current pretraining practices and evaluation protocols impose an effective ceiling on measured generalization.

Our evaluations point to one concrete contributor to this ceiling: split strategy. Homology-aware C3 splits (50) lowered and destabilized performance across the board, consistent with reduced homology leakage between training and evaluation data. KSMoFinder’s naive split yielded uniformly high performance with minimal model-to-model spread, whereas the C3-split version lowered overall scores and markedly increased variability. The same contrast emerged between Bernett and the SHS/STRING BFS PPI benchmarks, where most models scored consistently high, suggesting these datasets keep test examples closer to the training distribution. The pHopt case study provides further evidence: on the homology-aware rigor split, every MLP probe collapsed to zero or near-zero *R*^2^, against 0.393 for the best MLP probe on the standard split. Our multi-label subcellular compartment dataset points the same way. We deduplicated it at 80% identity and then split it randomly, so homologs below that threshold remain spread across train and test; every model scored far higher on it than on the clustersplit PPI benchmarks. Task difficulty may also contribute, since the subcellular and PPI datasets are not matched for label structure. Together, these patterns suggest that a meaningful share of the performance reported in pLM literature may reflect how well a model’s pretraining distribution overlaps with the evaluation set, rather than the depth and quality of its learned representations; homology-aware splitting and systematic quality controls such as cross-split deduplication should be adopted more broadly.

The pHopt case study illustrates how systematic bench-marking can reduce the need for extensive task-specific model development. The all-vs-all benchmark identified DPLM2_3B_’s representations as the strongest for this task, and a single transformer probe with sweep-optimized hyperparameters outperformed the EpHod Ensemble on standard regression metrics (*R*^2^ = 0.482 vs. 0.425; **Table 2**), despite EpHod’s far larger model search (42). Since EpHod, other pHopt predictors have been published, including CatOpt (*R*^2^ = 0.479) and the retrieval-enhanced Venus-DREAM (*R*^2^ = 0.509) (56, 57). Neither is directly comparable here. CatOpt merges EpHod’s training and validation sets, which EpHod had constructed by clustering at 20% sequence identity, then re-splits a random 10% for validation, removing the original homology-aware train/validation separation. Venus-DREAM retrieves labeled examples from the train set at inference and adapts on them rather than predicting from the sequence alone. We therefore keep EpHod as the matched comparison for this task. EpHod’s sample-weighted metrics, intended to emphasize sparse acidic and alkaline pH ranges through bin-inverse loss reweighting, remain higher than those of the transformer probe (**Supplementary Note 2**), since we applied no analogous rebalancing during probe training. We did not evaluate EpHod on the rigor split, so direct comparison under strict generalization conditions remains open.

The choice of adaptation strategy was also task-dependent in the two cases we tested. While the trans-former probe excelled on pHopt, a transfer learning comparison across six adaptation strategies on EC and GO-BP (**Supplementary Note 3**) showed no such advantage: the transformer probe offered negligible gains over the MLP probe on EC (MCC 0.806 vs. 0.800) and degraded performance on GO-BP (MCC 0.269 vs. 0.281), where LoRA achieved the highest score among all strategies tested. These contrasts reinforce the value of comparing multiple approaches under controlled, reproducible conditions before committing to expensive training.

Protify is an open-source platform for benchmarking protein language models across zero-shot, supervised, and transfer-learning settings, requiring minimal code. The need for easy, standardized evaluation will only grow as more and more pLMs and datasets are released. Protify was designed for that cycle, enabling the community to disentangle dataquality contributions from architectural design choices and training objectives. We expect it to connect pLM developers with life science domain scientists, making rigorous evaluation accessible regardless of computational expertise.

## Methods

### Framework Overview

Protify is an open-source platform for standardized workflows in pLM analysis. It implements an end-to-end evaluation loop comprising data loading, embedding extraction from pretrained pLMs, probe or fine-tuning training, evaluation, and reporting (**Fig. 1**). Models and datasets are accessed primarily via the Hugging Face Hub, with additional support for local CSV/FASTA files.

### Zero-shot Variant Effect Prediction

We used the ProteinGym substitutions benchmark to evaluate zero-shot performance for 23 pLMs. ProteinGym v1.0 was downloaded from `OATML-Markslab/ProteinGym_v1` with the Hugging Face datasets library on 22 August 2025, processing each assay into a separate .parquet file so that the full benchmark is never held in memory at once.

We used the masked-marginal scoring strategy described by Meier et al. (44). To avoid redundant computations, we grouped variants that share identical mutation indices and executed a single forward pass per unique set of masked positions, retrieving the log-probabilities for all substitutions at those positions. We scored a variant as the sum of the log-probabilities of the mutant amino acids at the masked positions minus the sum of the log-probabilities of the wild-type amino acids.

The original ProteinGym and E1 repositories instead mask a single residue at a time and cache the resulting log-its across variants. For single-residue substitutions, the three strategies are equivalent. However, for multi-site variants, the iterative approaches provide the wild-type identity at all non-masked mutation sites as context, which does not reflect the actual variant being scored. Simultaneous masking conditions each position only on the non-mutated context, avoiding information leakage from wild-type residues at co-mutated sites.

For sequences longer than a model’s context length (e.g., 1,024 tokens for ESM2), we used Tranception’s optimal windowing strategy (58), centering each window on the mean position of the mutated residues so that all mutation sites are included. The Tranception authors reported little difference between this and a sliding window.

The E1 models do not follow the pipeline described above. We scored them with the interface provided in (29), which handles masked-marginal scoring through the model’s native tokenizer, again substituting simultaneous for iterative masking. E1 supports an effective context length of 8,192 tokens, so we applied no windowing.

After scoring, we computed performance metrics with the ProteinGym DMS performance protocol and report the average Spearman rank correlation between model scores and experimental measurements for each assay (Table 1).

### Datasets

Protify supports 61 datasets directly, and users can pass any Hugging Face dataset with a sequence column and label column. We curated the native datasets from prior literature or directly from UniProt. Label descriptions, split sizes, and sources for each dataset are in **Tables S6** and **S7**.

### Models and embedding extraction

Protify supports 35 pretrained pLMs (**Table S8**), including ESM2, ESMC, ProtBert, ProtT5, ANKH, ANKH2, GLM2, DSM, DPLM, DPLM2, AMPLIFY, E1, and CaLM, as well as any custom pLM that loads with Hugging Face ‘*AutoModel*’ and outputs hidden states. Unless otherwise noted, we loaded models through the Hugging Face `transformers` library, with their original configurations and publicly released pretrained weights. For ESM2, ESMC, DPLM, DPLM2, DSM, ANKH, ANKH2, and E1, we used Synthyra’s FastPLMs implementations (59).

For each model, Protify provides an embedding wrapper class that returns per-residue embeddings from the frozen pretrained model, taken from the last hidden layer, as in prior work (48). For sequence-level representations, we computed the mean and variance across non-padding positions and concatenated the two pooled vectors.

CaLM_86M_ (25) is a codon-level BERT encoder pretrained with MLM on nucleotide codon triplets. Because all datasets except *ecoli_expression* contain amino acid sequences, Protify provides an `aa_to_dna` flag that deterministically reverse-translates each residue to its most frequent human synonymous codon. For *ecoli_expression*, which is already DNA, we scored CaLM_86M_ on the coding sequences directly and translated them to amino acids for the amino acid models.

To contextualize these pretrained models, we included three baseline controls, all pooled the same way as the pretrained embeddings:

- **Random features**: Each residue was assigned a fixed vector drawn from a random normal distribution. This negative control identifies dataset biases, such as la-bel imbalance, that might let a classifier score above chance without meaningful features.
- **One-Hot Encoding**: Each residue was represented as a one-hot vector over the amino acid alphabet. This baseline exposes only exact residue identity and composition, a simple proxy for sequence similarity or homology.
- **Random-Transformer**: a randomly initialized ESM2_35M_, acting as a fixed non-linear mapping over residue-level embeddings. Even with random token embeddings, attention makes similar sequence motifs attend to each other, so this approximates a sequence-homology control. Gains above it quantify what pretraining adds.

For multi-input datasets such as PPI prediction, Protify concatenates the vector or matrix embeddings of each named sequence column, padding as needed; the KSMoFinder datasets, for example, concatenate kinase, substrate, and motif embeddings.

### Supervised downstream evaluation

For the all-vs-all benchmark, we evaluated each pLM with a supervised probe trained on frozen embeddings, pooled as described above. Sequences were truncated to a maximum of 1,024 tokens to ensure a fair comparison across models with different native context lengths. For each dataset and model, we trained separate probes on the training splits, selected the best checkpoint by validation loss, and reported performance on the held-out test set. For KSMoFinder and soluprot, which were published without a validation split, we held out a random 10% of the training set for validation.

Protify’s MLP probe is a feed-forward network that maps pooled representations *x* ∈R^*d*^ to predictions *y* ∈R^*o*^, where *o* is the number of dataset labels/classes:

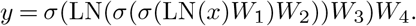

The weight matrices have shapes *W*_1_ ∈R^*d×h*^, *W*_2_ ∈ R^*h×h*^, *W*_3_ ∈R^*h×p*^, and *W*_4_ ∈ R^*p× o*^. We use ReLU activations *σ* (60) with hidden width *h* = 8,192. The intermediate width *p* is the smallest multiple of 256 that is at least 2*o*:

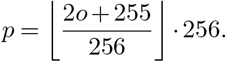

We applied dropout at rate 0.2 (61) immediately before each matrix multiplication except the first, and used no bias terms.

We trained all probes with the AdamW optimizer (62) under a cosine learning rate schedule, with a learning rate of 10_−4_, 100 warmup steps, a batch size of 64, and zero weight decay. Training ran for up to 200 epochs, with early stop-ping after three epochs of no improvement in validation loss. For datasets exceeding 350,000 training examples, evaluation was performed every 5,000 steps rather than per epoch.

We used cross-entropy loss for single-label classification, binary cross-entropy with logits loss for multi-label classification, and mean squared error loss for regression.

Protify computes metrics for each task type. For single-label classification, we report MCC, macro-F1, precision, recall, accuracy, AUROC, and AUPRC. For multi-label classification, we report MCC, Hamming loss, macro-averaged AUROC/AUPRC, and F1/precision/recall/accuracy after optimizing the decision threshold. For regression, we report Spearman *ρ*, Pearson *r* (with associated *p*-values), *R*^2^, MSE, MAE, and RMSE. In this work, we primarily report MCC for classification, as it remains informative under class imbalance while metrics such as F1 score and accuracy can produce misleadingly high scores (63), and *R*^2^ for regression, as it is more informative than alternative error metrics (64) and, unlike rank-based correlations, is sensitive to prediction scale and variance. We repeated experiments across three indepen-dent random seeds (seed, seed + 1, seed + 2) with seed = 42, and report the mean and standard deviation of the primary metric.

### Enzyme optimal pH case study

Gado et al. (42) introduced the pHopt dataset, 9,855 enzymes from BRENDA annotated with experimentally measured optimal pH values. We used the published train/validation/test splits (7,124/760/1,971) as-is. That splitting procedure first used a random 20% holdout and clustered only afterward, with MMseqs2 clustering at 20% sequence identity separating training from validation, so the test set had no homology control relative to the training data.

To evaluate without homology leakage, we built the `optimal_ph_rigor` dataset from BRENDA pHopt entries. We retrieved pH optimum annotations through the BRENDA SOAP API on April 16th, 2026, and mapped them to UniProt accessions to obtain canonical sequences. Where a sequence carried multiple pH entries, we used the average. We discarded sequences whose reported values spanned more than 1.0 pH unit or whose length fell outside 30 to 2,048 residues, then clustered the remainder with MMseqs2 at 30% sequence identity and 80% bidirectional coverage. We assigned entire clusters to train, validation, or test at a 90/5/5 ratio, stratified by integer pH bin to preserve label distribution across splits. We removed entries annotated as “assay at” in BRENDA, which give the pH at which the assay was run rather than the true catalytic optimum, from the validation and test sets. The resulting dataset contains 8,015 training, 430 validation, and 439 test sequences with zero sequence or cluster overlap between any pair of splits.

To reproduce EpHod’s published results, we evaluated their released predictions `(prediction.csv)` against the test split labels, matching by UniProt accession. We computed standard regression metrics through Protify, and sample-weighted regression metrics with bin-inverse weight-ing at bin borders [5.0, 9.0] to match EpHod’s evaluation protocol, using 100 bootstrap resamples for confidence intervals (**Fig. 3**).

Hyperparameters for the transformer probe were selected via Bayesian optimization through Protify’s Weights & Biases integration over 50 trials, about four hours of GH200 time. **Supplementary Table S9** gives the search space and the selected configuration. We trained the probe on perresidue embeddings from DPLM2_3B_ on both the standard and rigor splits.

## DATA AND RESOURCE AVAILABILITY

All datasets and models used in this study are publicly available on the Hugging Face Hub and are gathered in a single collection at huggingface.co/collections/GleghornLab/protify-paper-6ae9f64088657c668a562853, which includes the ProteinGym benchmark data and the trained optimal pH probes. Tables S6 and S7 describe each dataset and its split sizes, and Table S8 lists the pretrained models. The optimal pH dataset with the original EpHod splits is available at GleghornLab/optimal_ph, and the homology-aware rigor split at `GleghornLab/optimal_ph_rigor`. Protify is freely available at github.com/Gleghorn-Lab/Protify.

## ACKNOWLEDGEMENTS

The authors thank Katherine M. Nelson, Ph.D., for reviewing and commenting on drafts of the manuscript. This work was partly supported by the University of Delaware Graduate College through the Unidel Distinguished Graduate Scholar Award (LH), the National Science Foundation through ART 2331440 (LH) and NAIRR pilot 240064 (JPG), and the National Institutes of Health through NIGMS T32GM142603 (LH, NR) and R01HL178817 (JPG).

## AUTHOR CONTRIBUTIONS

Conceived and designed the experiments (NR, LH, JPG), per-formed the experiments (NR, LH, TP), contributed code (NR, LH, TP, CH, LHav), analyzed the data (NR, LH, TP, CH), wrote the paper (NR, LH, TP, JPG), supervised the work (LH, JPG), and acquired funding (LH, JPG).

## Supplementary Information

### Supplementary Note 1: Pooling strategy selection

To identify the best-performing embedding pooling strategy for the supervised vector benchmark, we ran a preliminary study using ESMC300M embeddings on a subset of the datasets. We evaluated all individual pooling operations (mean, max, min, *p* = 2 norm, median, standard deviation, var, cls (first token), eos (last non-pad token)), as well as all pairwise combinations formed by concatenating two pooling types. We used the optimal-temp, optimal-ph, fitness-pred, enzyme-*kcat*, stability-pred, MB, DeepLoc-2, DeepLoc-10, cloning-clf, and mat-production datasets. For each pooling configuration, we trained a Random Forest on the pooled embeddings, a classifier (n_estimators = 100, max_depth = 30, log-loss criterion) for classification tasks, and a regressor (max_depth = 100, no bootstrap) for regression tasks, both with random_state = 42. Sequences were truncated to 2,048 residues. For pairwise pooling combinations, we evaluated concatenation, alongside single-pooling baselines. We then ranked each configuration on every dataset by its primary metric (MCC for classification, the average of Pearson correlation coefficient and R^2^ for regression) and summed the ranks across datasets to obtain a total rank score. Lower scores indicate more consistent performance.

**Table S1.** Pooling strategies ranked by their summed per-dataset rank across 10 datasets, using ESMC300M embeddings and a random forest (lower is better). Pairwise strategies concatenate the two pooled representations (⊕ ).

| Ranks 1-23 |  |  | Ranks 24-45 |  |  |
|---|---|---|---|---|---|
| Rank | Strategy | Score | Rank | Strategy | Score |
| 1 | **mean ⊕ var** | **60** | 24 | cls ⊕ eos | 228 |
| 2 | **mean ⊕ std** | **91** | 25 | min ⊕ std | 232 |
| 3 | **median ⊕ var** | **101** | 26 | min ⊕ var | 234 |
| 4 | mean ⊕ median | 104 | 27 | max ⊕ var | 239 |
| 5 | mean ⊕ cls | 116 | 28 | var | 246 |
| 6 | median ⊕ std | 118 | 29 | min ⊕ eos | 248 |
| 7 | mean | 119 | 30 | norm ⊕ cls | 251 |
| 8 | median ⊕ cls | 119 | 31 | max ⊕ std | 251 |
| 9 | mean ⊕ min | 131 | 32 | var ⊕ eos | 252 |
| 10 | mean ⊕ max | 142 | 33 | std ⊕ eos | 254 |
| 11 | eos | 144 | 34 | cls | 254 |
| 12 | median | 144 | 35 | std | 256 |
| 13 | mean ⊕ eos | 152 | 36 | min ⊕ norm | 264 |
| 14 | min ⊕ median | 157 | 37 | max ⊕ eos | 268 |
| 15 | median ⊕ eos | 163 | 38 | max ⊕ min | 274 |
| 16 | max ⊕ median | 176 | 39 | max ⊕ norm | 283 |
| 17 | mean ⊕ norm | 177 | 40 | norm | 286 |
| 18 | var ⊕ cls | 186 | 41 | norm ⊕ std | 298 |
| 19 | min ⊕ cls | 196 | 42 | max | 300 |
| 20 | norm ⊕ median | 199 | 43 | norm ⊕ var | 302 |
| 21 | std ⊕ cls | 203 | 44 | norm ⊕ eos | 325 |
| 22 | min | 212 | 45 | std ⊕ var | 328 |
| 23 | max ⊕ cls | 222 |  |  |  |

Table S1 reports the total rank score for all 45 pooling configurations tested. Mean-variance concatenation achieved the lowest total rank score (60), the most consistent performance across datasets. This was followed by mean-std (91) and median-var (101). Based on these results, we adopted mean-variance pooling as the default strategy for the vector benchmark experiments and Protify itself. Full code and results are available at https://github.com/Gleghorn-Lab/Pooling-PLM-Hidden-States.

### Supplementary Note 2: Enzyme optimal pH: sample-weighted evaluation

Table 2 reports standard regression metrics, on which the DPLM23B transformer probe ranks above EpHod. However, EpHod’s evaluation protocol also includes sample-weighted metrics designed to emphasize performance on under-represented pH ranges (42). Following their protocol, we computed bin-inverse-weighted metrics with bin borders at pH 5.0 and 9.0, partitioning predictions into acidic (*<*5.0), neutral (5.0-9.0) and alkaline (*>*9.0) bins. Each bin’s contribution is weighted inversely to its frequency, so that sparse acidic and alkaline predictions have a greater influence on the aggregate score. We obtained confidence intervals from 100 bootstrap resamples.

On sample-weighted metrics, the EpHod Ensemble outperforms the transformer probe (Table S2): it achieves *R*^2^ = 0.681 versus 0.600, weighted Pearson *r* = 0.843 versus 0.831, and weighted RMSE = 1.274 versus 1.426. This reversal was largely expected, as EpHod was trained with a bin-inverse loss function that explicitly penalizes errors on under-represented pH ranges, whereas the transformer probe achieves F1 scores of 0.651, 0.519, and 0.000 on the acidic, neutral, and alkaline bins, respectively, indicating that it does not recover signal for the rarest pH range. Incorporating loss reweighting into probe training would likely close the gap on sample-weighted metrics while preserving the probe’s advantage on standard metrics.

**Table S2.**
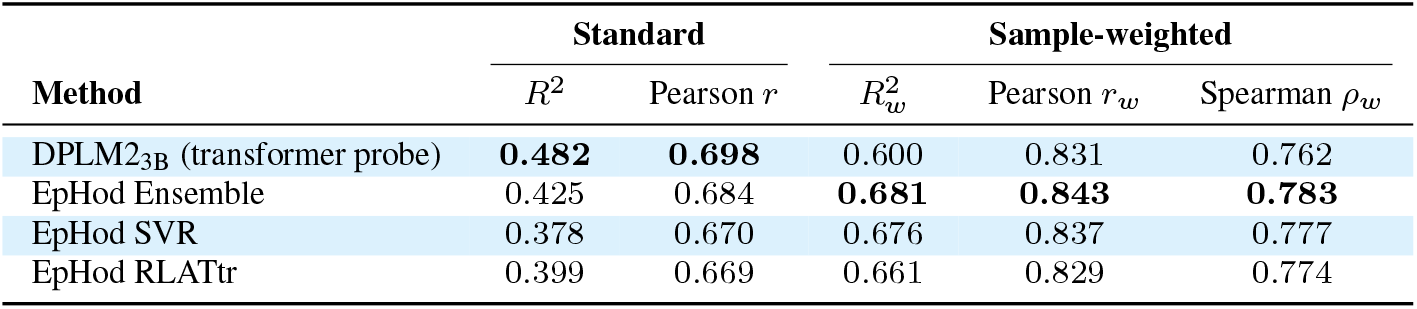
Standard versus sample-weighted regression metrics on the pHopt test set (*n*=1,971). Sample-weighted metrics use bininverse weighting with bin borders [5.0, 9.0] and 100 bootstrap resamples, following EpHod’s evaluation protocol (42). Bold indicates best performance per column. The transformer probe ranks first on standard metrics but falls below the EpHod Ensemble on sample-weighted metrics, reflecting the absence of bin-inverse loss reweighting during probe training.

|  | **Standard** |  | **Sample-weighted** |  |  |
|---|---|---|---|---|---|
| **Method** | *R*<sup>2</sup> | Pearson *r* | *R*<sub>*w*</sub><sup>2</sup> | Pearson *r*<sub>*w*</sub> | Spearman *ρ*<sub>*w*</sub> |
| DPLM2<sub>3B</sub> (transformer probe) | **0.482** | **0.698** | 0.600 | 0.831 | 0.762 |
| EpHod Ensemble | 0.425 | 0.684 | **0.681** | **0.843** | **0.783** |
| EpHod SVR | 0.378 | 0.670 | 0.676 | 0.837 | 0.777 |
| EpHod RLATtr | 0.399 | 0.669 | 0.661 | 0.829 | 0.774 |

### Supplementary Note 3: Transfer learning on difficult tasks

For complex, high-dimensional annotation tasks, the choice of adaptation strategy can be just as critical as the choice of pretrained model. This is particularly evident in sparse multi-label datasets, such as Enzyme Commission (EC) number prediction and Gene Ontology Biological Process (GO-BP) annotation, where label spaces contain hundreds to thousands of categories, and training data are limited. In these scenarios, an MLP probe, while computationally efficient, may lack the expressivity to capture non-linear relationships between the embedding and label spaces, potentially sacrificing performance.

To compare adaptation strategies, we evaluated six approaches on the EC and GO-BP datasets using ESMC600M. All six strategies shared identical early stopping criteria (a patience of three epochs on validation loss), a cosine learning rate schedule with a learning rate of 10_*−*4_, a maximum of 200 training epochs, and three independent random seeds, with the initial seed being 1,184,471,526 (automatically generated from the current time). Unless otherwise noted, hyperparameters match the MLP probe training protocol described above.

**Fig. S1.**
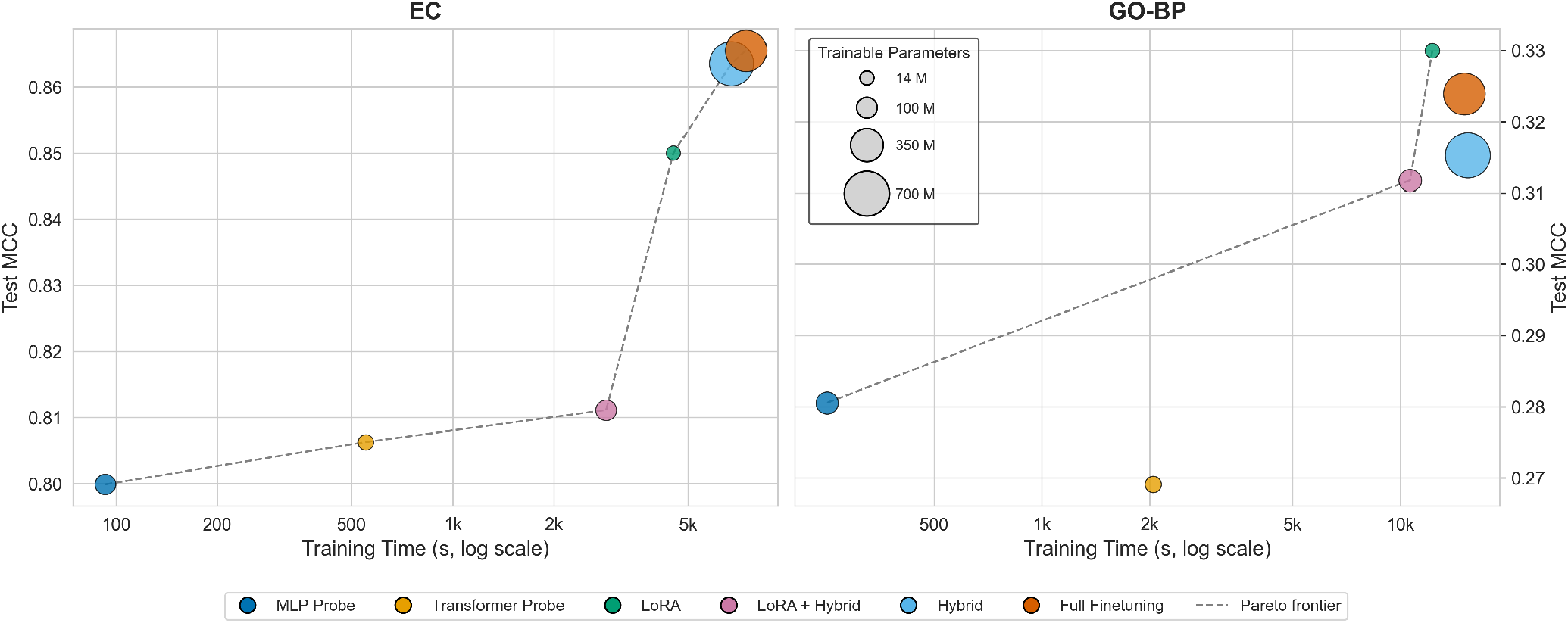
Efficiency-performance trade-offs for adaptation strategies on sparse multi-label annotation. Pareto analysis of six transfer learning strategies using ESMC600M on EC (left) and GO-BP (right). The x-axis shows total training time in seconds (log scale) required to reach the early stopping patience under a fixed training protocol. The y-axis shows test-set MCC (mean over three random seeds). Marker size indicates the number of trainable parameters for each strategy. The dashed line denotes the Pareto frontier (strategies not dominated in both training time and MCC).

On the EC dataset, performance tracked computational cost almost monotonically (**Fig. S1, Table S3**). The MLP probe achieved an MCC of 0.800 in just 93 seconds, while full finetuning reached 0.866 after 7,423 seconds, a roughly 8% increase in performance at an 80-fold increase in training time. Notably, LoRA emerged as a compelling strategy on this task, achieving an MCC of 0.850 at 4,527 seconds, capturing 76% of the performance gain of full finetuning at 61% of the training cost. The transformer probe achieved an MCC of 0.806, only 0.006 higher than the MLP probe despite a six-fold increase in training time, suggesting the bottleneck lies in the frozen representation rather than the probe architecture. The LoRA + Hybrid strategy (0.811) underperformed both the standalone LoRA and Hybrid (0.864) strategies, indicating that naively combining adaptation strategies does not guarantee higher gains.

On the GO-BP dataset, the efficiency-performance landscape changed substantially (**Fig. S1**). Most strikingly, LoRA achieved the highest MCC of all strategies (0.330), surpassing both full finetuning (0.324) and the Hybrid approach (0.315). This shift suggests that in sparser, higher-dimensional label spaces, the implicit regularization from low-rank constraints can help prevent the model from overfitting to limited training signal. The MLP probe remained a strong baseline (0.281), while the transformer probe performed worse (0.269). The LoRA + Hybrid strategy (0.312) again underperformed LoRA alone, reinforcing that the hybrid approach’s two-phase training may be counterproductive when combined with LoRA.

The Pareto frontiers across both tasks highlight the MLP probe and LoRA as the most consistently efficient adaptation strategies in both datasets, with the former offering competitive performance with minimal overhead and the latter delivering substantial gains while remaining computationally tractable. Full finetuning and the hybrid approaches yielded the highest scores on EC, but at substantially greater cost, and neither was able to surpass LoRA on GO-BP.

**Table S3.**
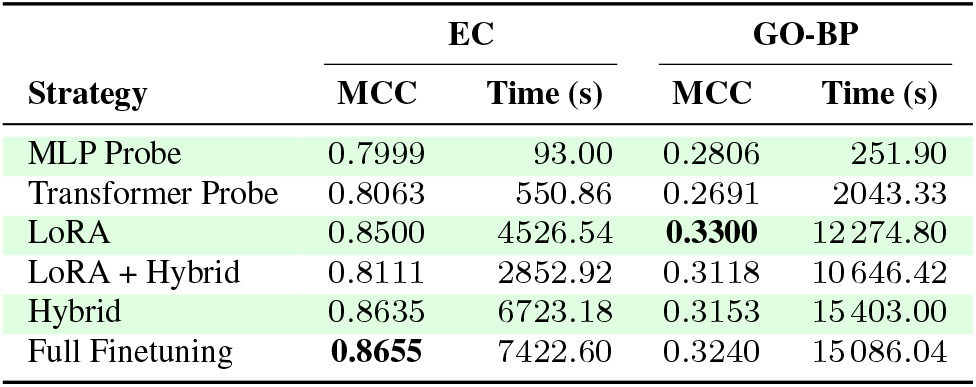
Transfer learning strategy comparison on EC and GO-BP datasets using ESMC_600M_. MCC (mean over three random seeds) and total training time are reported. Bold indicates the best MCC per dataset.

| Strategy | EC |  | GO-BP |  |
|---|---|---|---|---|
|  | MCC | Time (s) | MCC | Time (s) |
| MLP Probe | 0.7999 | 93.00 | 0.2806 | 251.90 |
| Transformer Probe | 0.8063 | 550.86 | 0.2691 | 2043.33 |
| LoRA | 0.8500 | 4526.54 | **0.3300** | 12 274.80 |
| LoRA + Hybrid | 0.8111 | 2852.92 | 0.3118 | 10 646.42 |
| Hybrid | 0.8635 | 6723.18 | 0.3153 | 15 403.00 |
| Full Finetuning | **0.8655** | 7422.60 | 0.3240 | 15 086.04 |

We note that these results are not intended as a direct comparison to published state-of-the-art scores on EC and GO-BP. For reference, FusionProt (65) reports Fmax scores of 0.877 on EC and 0.345 on GO-BP for an ESM2-650 baseline, which fall in a similar range to our MCC scores for full finetuning of 0.866 and 0.324, respectively. However, the two metrics measure different quantities (Fmax selects the optimal prediction threshold per protein, while MCC captures all four confusion matrix categories), and the underlying dataset splits are likely different as well. Our splits come directly from SaProt, which used a 30% sequence-identity filter, while FusionProt applies a similar Needleman-Wunsch 30% sequence-identity filter on top of TorchDrug’s default splits, though their exact splits are not publicly available. Given these differences, this case study does not compare performance against external baselines, but compares adaptation strategies under identical conditions so observed differences reflect the strategy itself.

On EC, where the label space is smaller with more training examples per class, full finetuning and the hybrid approach yielded the highest absolute performance, whereas on GO-BP, where labels are spread across a larger and sparser ontology, LoRA achieved the highest MCC of all strategies, surpassing full finetuning. This reversal suggests that low-rank adaptation constraints may be advantageous when per-class training signal is limited, likely preventing overfitting to noise. Despite being the simplest and least expensive strategy, the MLP probe recovered the majority of the performance achieved by more complex methods on both datasets, occupying the Pareto frontier alongside LoRA. This indicates that much of the task-relevant signal is already accessible in frozen representations, and that researchers can capture most downstream performance at a fraction of the cost before deciding whether further adaptation is warranted.

We evaluated all six strategies under a single default training recipe without per-dataset hyperparameter tuning. On sparse, high-dimensional tasks like GO-BP, strategies that update model weights are known to be sensitive to learning rate, warmup, and regularization choices, and their absolute performance would likely improve with dataset-specific tuning. However, in compute-constrained settings where per-task hyperparameter sweeps are not feasible, the MLP probe serves as a reliable and practical default under a fixed training protocol.

#### Transformer probe

The transformer probe operates on per-residue embeddings *E* ∈ R^*l×d*^ rather than pooled vectors. An input projec-tion

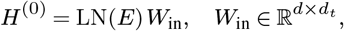

mapping inputs to a hidden dimension of *d*_*t*_ = 1,024. This was followed by *n* = 2 transformer encoder blocks with RoPE (66, 67), with SwiGLU MLP (68). Each transformer block used PyTorch LayerNorm before the queries and keys, four attention heads, and an expansion ratio of 8/3, rounded to the nearest multiple of 256 in the MLP. After the transformer layer, mean-variance pooling is applied over non-padding positions and concatenated:

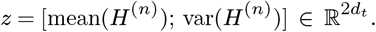

A classifier MLP produces the final output:

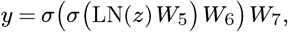

Where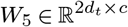, *W*_6_ ∈ R^*c×p*^, *W*_7_ ∈R^*p×o*^, and *σ* is ReLU. *c* = 1,024, with *p* computed identically to the MLP probe. We applied 10% dropout in the transformer block and 20% in the classifier. Bias terms were not used. We trained the probe with a batch size of 64 and 100 warmup steps.

#### Full finetuning

In the full finetuning strategy, all parameters of the pretrained pLM were unfrozen and trained end-to-end with the model’s native classification head as provided by each respective implementation. We used a batch size of 8 with gradient accumulation over 4 steps (effective batch size of 32), and 1,000 warmup steps.

#### LoRA

For the LoRA strategy (69), the pretrained weights *W*_0_ are frozen, and each weight in the attention layer is decomposed into:

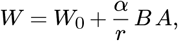

Where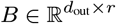 and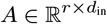 are the only trainable parameters. LoRA adapters were applied to the query, key, value, and output projection matrices in every attention layer, with rank *r* = 8, scaling factor *α* = 32, and 1% LoRA dropout. We used a batch size of 16 with gradient accumulation over 2 steps, and 1,000 warmup steps.

#### Hybrid probe

The hybrid strategy consists of a two-phase training procedure. In phase 1, an MLP probe (as described above) is trained on frozen pooled embeddings. In phase 2, the trained probe is attached to the base pLM and the composite model is finetuned end-to-end:

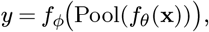

where *f*_*ϕ*_ denotes the MLP probe, *f*_*θ*_ the pLM, and Pool the concatenated mean-variance pooled features. The total training time reported is the sum of both phases. Phase 1 uses the MLP probe training protocol, and phase 2 the full finetuning protocol.

#### LoRA-hybrid

This strategy combines LoRA with the two-phase hybrid approach. Phase 1 trains an MLP probe on frozen embeddings, and in phase 2 LoRA adapters (*B, A* matrices) are inserted into the attention layers of the base model. The composite model is finetuned end-to-end, with only the LoRA adapters and probe parameters being updated during phase 2, while the pretrained weights *W*_0_ remain frozen. The training protocol and LoRA configuration match the ones described above.

### Supplementary Note 4: Vector benchmark heatmap views

**Figure S2** shows the same all-vs-all heatmap as Fig. 2, but includes standard deviations across three random seeds alongside the mean scores, showing seed-to-seed variability for each model-dataset combination. **Figure S3** shows the normalized all-vs-all heatmap, where each dataset’s scores are min-max scaled to [0, 1] so the worst-performing model receives 0 and the best receives 1. This view allows a fair comparison of relative model rankings across all 32 datasets by ignoring absolute scale differences between tasks.

**Fig. S2.**
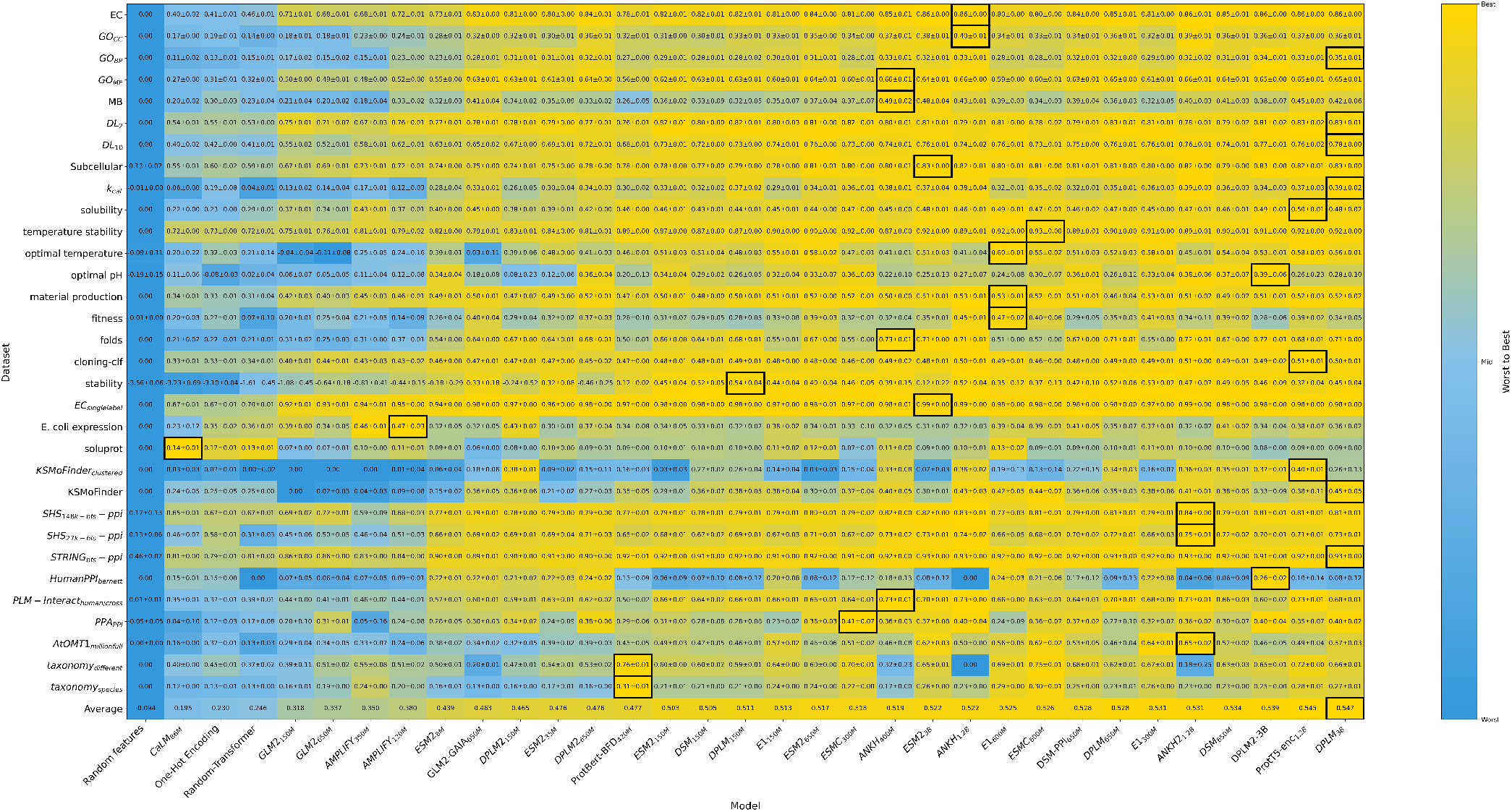
All-vs-all supervised vector benchmark with standard deviations. Heatmap of test set performance for each modeldataset pair, identical to Fig. 2 but with standard deviations across three random seeds shown alongside the mean score.

**Fig. S3.**
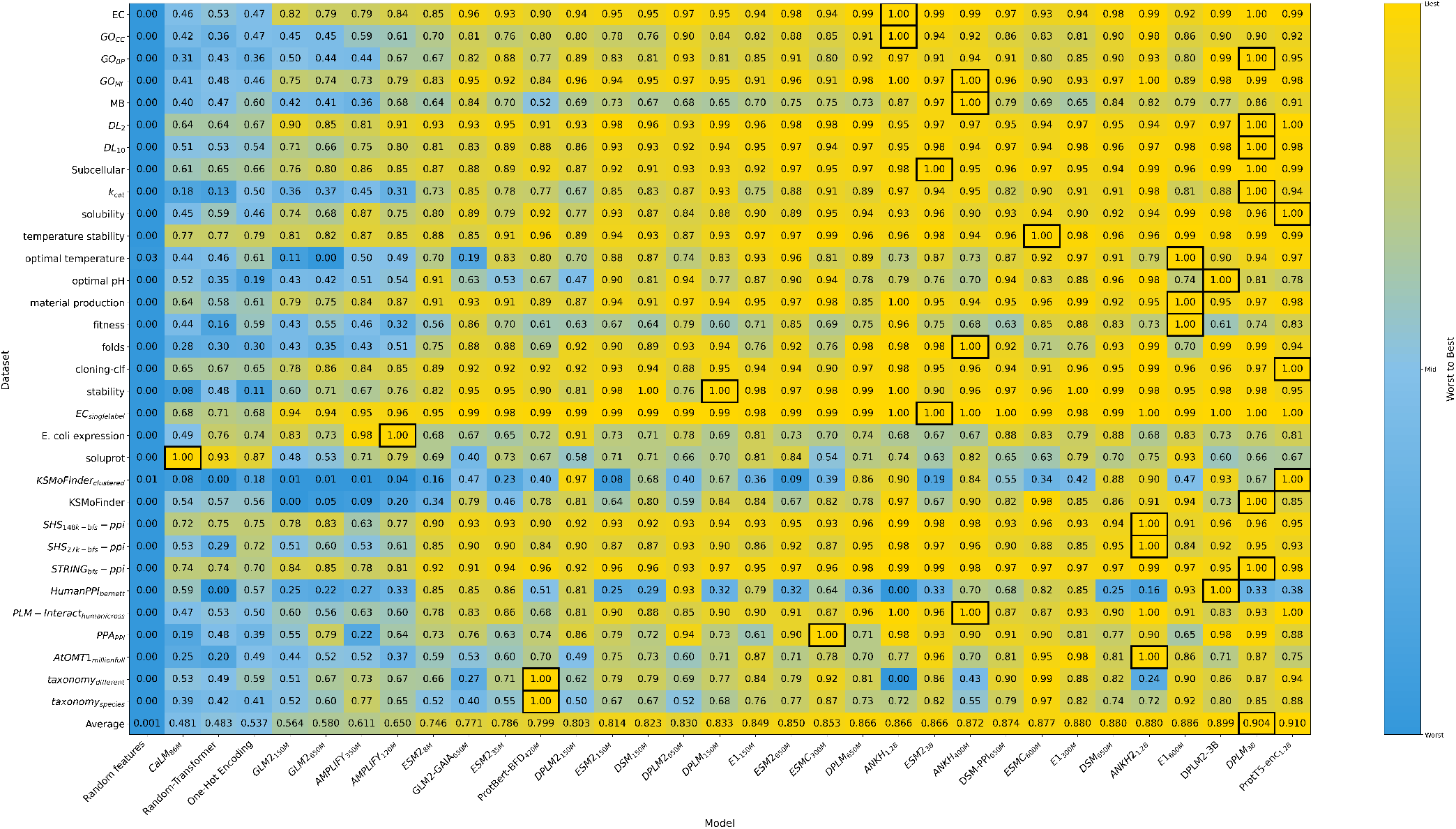
Normalized all-vs-all supervised vector benchmark. Each dataset’s scores are min-max normalized to [0, 1] so that the lowest-scoring model maps to 0 (blue) and the highest-scoring model maps to 1 (yellow). The average row reports the mean normalized score across all 32 datasets for each model. Models are sorted by this average in ascending order from left to right. Black outlines highlight the best performer per dataset.

### Supplementary Note 5: Winner margin analysis

Table S4 reports, for each of the 32 vector benchmark datasets, the best- and second-best performing models by mean test-set score across three seeds, the margin Δ between them, and the Benjamini-Hochberg adjusted *p*-value from a two-sided Welch’s *t*-test. Before correction, 31 of 32 raw *p*-values exceed 0.05, and after correction, none reach statistical significance at *α* = 0.05.

These results should be interpreted with caution. With only *n* = 3 seeds per model, the Welch tests are severely underpowered (hence the high *p*-values), so non-significance is largely expected for the small margins observed, and does not imply equivalency. Additionally, because all models share the same three random seeds, a paired test would be more appropriate; however, only summary statistics were retained. The table is therefore best read as a descriptive summary showing how thin the margins between top models are relative to run-to-run variability, and not as a formal test of equivalence.

### Supplementary Note 6: Scikit-learn case study

We implemented a supervised learning pipeline using scikit-learn to evaluate the predictive capacity of protein language model embeddings for PPI prediction. We screened 10 classifiers supported by LazyPredict (70): *LogisticRegression, RandomForestClassifier, XGBClassifier, LGBMClassifier, BernoulliNB, GaussianNB, NearestCentroid, PassiveAggressiveClassifier, RidgeClassifier*, and *DummyClassifier*. We trained models on concatenated mean- and variance-pooled ESMC600M embeddings (size 4,608 total) using the gold-standard PPI dataset (52, 53), partitioned into training (n=163,192), validation (n=59,260), and test (n=52,048) sets. After the initial screen, we optimized the top-performing classifier using *RandomizedSearchCV* with 100 iterations of 3-fold cross-validation. The optimization hyperparameters included `n_estimators` (100–500 in steps of 100), `max_depth` (3–8), `learning_rate` (0.01–0.2), `subsample` (0.6–1.0), and `colsample_bytree` (0.6–1.0). We evaluated the final optimized *XGBClassifier* on the held-out test set to obtain unbiased performance estimates.

An initial screen of ten scikit-learn classifiers on the Bernett PPI dataset revealed that XGBoost (*XGBClassifier*) achieved the highest MCC (Table S5). Most classifiers produced MCC scores at or near zero, indicating no meaningful predictive ability beyond random chance. Hyperparameter optimization of the best-performing *XGBClassifier* via *RandomizedSearchCV* (100 iterations, 3-fold cross-validation) yielded a best cross-validation MCC of 0.197. With the configuration (`n_estimators=500`, `max_depth=7`, `learning_rate=0.1`, `subsample=1.0`, and `colsample_bytree=1.0`), *XGBClassifier* achieved a test-set MCC of 0.17. This score is consistent with the uniformly low performance observed across most pLMs on this C3-split dataset in the vector benchmark, and is only slightly lower than 0.21 achieved by the MLP probe.

**Table S4.** Winner-margin analysis for the 32 datasets. For each dataset, the best and second-best performing models are shown with their mean score ± standard deviation across three random seeds (*n* = 3). Δ is the difference between the two means; *p*_adj_ is the two-sided Welch’s *t*-test *p*-value after Benjamini-Hochberg correction across all 32 comparisons. MCC is used for classification tasks; *R*^2^ for regression tasks.

| **Dataset** | **Best Model** | **Score** | **2nd Best** | **Score** | **Δ** | *p*<sub>**adj**</sub> |
|---|---|---|---|---|---|---|
| *Classification (MCC)* |  |  |  |  |  |  |
| GO-BP | DPLM<sub>3B</sub> | 0.3463 ± 0.0124 | DPLM2<sub>3B</sub> | 0.3439 ± 0.0107 | 0.0024 | 0.958 |
| GO-CC | ANKH<sub>1.2B</sub> | 0.3985 ± 0.0052 | ANKH2<sub>1.2B</sub> | 0.3902 ± 0.0070 | 0.0083 | 0.877 |
| DeepLoc-10 | DPLM<sub>3B</sub> | 0.7793 ± 0.0025 | DPLM2<sub>3B</sub> | 0.7653 ± 0.0148 | 0.0140 | 0.877 |
| DeepLoc-2 | DPLM<sub>3B</sub> | 0.8317 ± 0.0092 | ProtT5<sub>enc-1.2B</sub> | 0.8311 ± 0.0154 | 0.0006 | 0.959 |
| EC | ANKH<sub>1.2B</sub> | 0.8642 ± 0.0022 | DPLM<sub>3B</sub> | 0.8599 ± 0.0016 | 0.0043 | 0.877 |
| KSMoFinder | DPLM<sub>3B</sub> | 0.4476 ± 0.0519 | ESMC<sub>600M</sub> | 0.4402 ± 0.0700 | 0.0074 | 0.958 |
| MB | ANKH<sub>400M</sub> | 0.4934 ± 0.0157 | ESM2<sub>3B</sub> | 0.4776 ± 0.0367 | 0.0159 | 0.947 |
| GO-MF | ANKH<sub>400M</sub> | 0.6611 ± 0.0065 | ANKH<sub>1.2B</sub> | 0.6600 ± 0.0026 | 0.0011 | 0.958 |
| SL13 | ESM2<sub>3B</sub> | 0.8318 ± 0.0032 | DPLM<sub>3B</sub> | 0.8307 ± 0.0041 | 0.0010 | 0.958 |
| gold-ppi | DPLM2<sub>3B</sub> | 0.2574 ± 0.0214 | DPLM2<sub>650M</sub> | 0.2405 ± 0.0180 | 0.0169 | 0.877 |
| cloning-clf | ProtT5<sub>enc-1.2B</sub> | 0.5111 ± 0.0072 | ANKH2<sub>1.2B</sub> | 0.5084 ± 0.0044 | 0.0027 | 0.947 |
| taxonomy_different | ProtBert-BFD<sub>420M</sub> | 0.7609 ± 0.0127 | ESMC<sub>600M</sub> | 0.7523 ± 0.0150 | 0.0087 | 0.947 |
| ec-active | ESM2<sub>3B</sub> | 0.9869 ± 0.0013 | ANKH<sub>1.2B</sub> | 0.9852 ± 0.0047 | 0.0017 | 0.947 |
| ecoli_expression | AMPLIFY<sub>120M</sub> | 0.4691 ± 0.0308 | AMPLIFY<sub>350M</sub> | 0.4580 ± 0.0096 | 0.0110 | 0.947 |
| number-of-folds | ANKH<sub>400M</sub> | 0.7253 ± 0.0091 | ANKH2<sub>1.2B</sub> | 0.7200 ± 0.0063 | 0.0053 | 0.947 |
| KSMoFinder-clustered | ProtT5<sub>enc-1.2B</sub> | 0.3951 ± 0.0056 | DPLM2<sub>150M</sub> | 0.3814 ± 0.0139 | 0.0137 | 0.877 |
| mat-production | E1<sub>600M</sub> | 0.5345 ± 0.0051 | ANKH<sub>1.2B</sub> | 0.5321 ± 0.0102 | 0.0024 | 0.958 |
| plm-interact | ANKH<sub>400M</sub> | 0.7285 ± 0.0069 | ANKH2<sub>1.2B</sub> | 0.7278 ± 0.0098 | 0.0007 | 0.958 |
| shs148-ppi-bfs | ANKH2<sub>1.2B</sub> | 0.8352 ± 0.0024 | ANKH<sub>1.2B</sub> | 0.8305 ± 0.0051 | 0.0047 | 0.877 |
| shs27-ppi-bfs | ANKH2<sub>1.2B</sub> | 0.7544 ± 0.0114 | ANKH<sub>1.2B</sub> | 0.7447 ± 0.0166 | 0.0098 | 0.947 |
| string-ppi-bfs | DPLM<sub>3B</sub> | 0.9335 ± 0.0011 | ANKH2<sub>1.2B</sub> | 0.9302 ± 0.0027 | 0.0032 | 0.877 |
| solubility | ProtT5<sub>enc-1.2B</sub> | 0.4976 ± 0.0084 | E1<sub>600M</sub> | 0.4902 ± 0.0054 | 0.0074 | 0.877 |
| soluprot | CaLM<sub>86M</sub> | 0.1375 ± 0.0134 | E1<sub>600M</sub> | 0.1282 ± 0.0174 | 0.0093 | 0.947 |
| taxonomy_species | ProtBert-BFD<sub>420M</sub> | 0.3140 ± 0.0107 | ESMC<sub>600M</sub> | 0.3035 ± 0.0061 | 0.0106 | 0.877 |
| temp-stability | ESMC<sub>600M</sub> | 0.9302 ± 0.0023 | E1<sub>600M</sub> | 0.9230 ± 0.0014 | 0.0073 | 0.468 |
| *Regression (R<sup>2</sup>)* |  |  |  |  |  |  |
| enzyme-*k*<sub>*cat*</sub> | DPLM<sub>3B</sub> | 0.3941 ± 0.0152 | ANKH2<sub>1.2B</sub> | 0.3877 ± 0.0329 | 0.0063 | 0.958 |
| fitness-prediction | E1<sub>600M</sub> | 0.4659 ± 0.0229 | ANKH<sub>1.2B</sub> | 0.4465 ± 0.0130 | 0.0195 | 0.877 |
| million_full | ANKH2<sub>1.2B</sub> | 0.6491 ± 0.0157 | E1<sub>300M</sub> | 0.6374 ± 0.0089 | 0.0117 | 0.877 |
| optimal-ph | DPLM2<sub>3B</sub> | 0.3931 ± 0.0631 | ANKH2<sub>1.2B</sub> | 0.3839 ± 0.0638 | 0.0092 | 0.958 |
| optimal-temperature | E1<sub>600M</sub> | 0.6014 ± 0.0112 | ProtT5<sub>enc-1.2B</sub> | 0.5827 ± 0.0266 | 0.0188 | 0.877 |
| PPA-ppi | ESMC<sub>300M</sub> | 0.4056 ± 0.0668 | DPLM<sub>3B</sub> | 0.4008 ± 0.0164 | 0.0048 | 0.958 |
| stability-pred | DPLM<sub>150M</sub> | 0.5377 ± 0.0401 | E1<sub>300M</sub> | 0.5266 ± 0.0243 | 0.0111 | 0.958 |

**Total:** 32 datasets. Significant (*p* < 0.05): 0/32 (0%)

**Table S5.** Initial classifier screening on the Bernett PPI dataset using concatenated mean-variance-pooled ESMC600M embeddings. MCC is reported on the validation set. Classifiers are sorted by MCC in descending order.

| **Classifier** | **MCC** |
|---|---|
| XGBClassifier | 0.10 |
| LGBMClassifier | 0.07 |
| RandomForestClassifier | 0.05 |
| LogisticRegression | 0.01 |
| BernoulliNB | 0.01 |
| NearestCentroid | 0.00 |
| RidgeClassifier | 0.00 |
| GaussianNB | 0.00 |
| DummyClassifier | 0.00 |
| PassiveAggressiveClassifier | 0.00 |

### Supplementary Note 7: Supported datasets and models

Tables S6 and S7 list the 61 curated datasets in Protify: the 32 used in the vector benchmark and 29 others. Table S8 lists the supported pretrained models.

**Table S6.** Vector benchmark datasets used in the comprehensive comparison. **Legend:** Multi-Label datasets are shown in ; Multi-Class in ; Binary Classification in ; and Regression in .

| Dataset | Description | Multi-Input | Size (Splits) | Source |
|---|---|---|---|---|
| **EC** | Enzyme function prediction using EC numbers.<br>*Labels:* 585 EC Categories | No | *Train:* 13,090<br>*Val:* 1,465<br>*Test:* 1,604 | (71) |
| **GO-CC** | Gene Ontology Cellular Component (GO-CC) prediction.<br>*Labels:* 320 Terms | No | *Train:* 26,225<br>*Val:* 2,904<br>*Test:* 3,350 | (71) |
| **GO-BP** | Gene Ontology Biological Process (GO-BP) prediction.<br>*Labels:* 1,943 Terms | No | *Train:* 26,225<br>*Val:* 2,904<br>*Test:* 3,350 | (71) |
| **GO-MF** | Gene Ontology Molecular Function (GO-MF) prediction.<br>*Labels:* 489 Terms | No | *Train:* 26,225<br>*Val:* 2,904<br>*Test:* 3,350 | (71) |
| **SL13** | Custom UniProt subcellular localization dataset from experimentally annotated entries (Query: 2025-06-17). Filtered for length 20-2048, clustered at 80% identity (CD-HIT, *n*=5), keeping representative sequences. Multi-label binary vector over 13 locations: Cytoplasm, Nucleus, Secreted, Membrane, Mitochondria, Plastid, ER, Vacuole, Lysosome, Golgi Apparatus, Peroxisome, Chloroplast, Cell Wall. Deduplicated and randomly split with 1k validation and 1k test.<br>*Labels:* 13 Locations | No | *Train:* 32,638<br>*Val:* 1,000<br>*Test:* 1,000 | (19) |
| **shs27-ppi-bfs** | SHS27k PPI (CD-HIT 40%, BFS splits).<br>*Labels:* 7 Types (PTM, Reaction, Catalysis, Inhibition, Binding, Activation, Expression) | Yes | *Train:* 4,440<br>*Val:* 1,480<br>*Test:* 1,481 | (72–74) |
| **shs148-ppi-bfs** | SHS148k PPI (CD-HIT 40%, BFS splits).<br>*Labels:* 7 Types (PTM, Reaction, Catalysis, Inhibition, Binding, Activation, Expression) | Yes | *Train:* 26,038<br>*Val:* 8,679<br>*Test:* 8,680 | (72–74) |
| **string-ppi-bfs** | STRING-derived PPI (CD-HIT 40%, BFS splits).<br>*Labels:* 7 Types (PTM, Reaction, Catalysis, Inhibition, Binding, Activation, Expression) | Yes | *Train:* 343,540<br>*Val:* 114,514<br>*Test:* 114,514 | (72, 74, 75) |
| **DeepLoc-10** | Protein subcellular localization classification.<br>*Labels:* 10 Locations | No | *Train:* 8,743<br>*Val:* 2,190<br>*Test:* 2,745 | (71) |
| **ec-active** | Custom UniProt EC dataset with active site info from entries with manual GO assertions (Query: 2025-05-30). Entries with ambiguous (‘-’) or multiple EC numbers removed. Filtered for length 20-2048, clustered at 80% identity (CD-HIT, *n*=3), keeping representative sequences. EC classes with ≥100 examples retained (260 classes). Deduplicated, favoring entries with active-site annotations. 13.8% of training set has active-site info. Randomly split with 1k validation and 1k test (both exclusively active-site annotated).<br>*Labels:* 260 EC Categories | No | *Train:* 80,212<br>*Val:* 1,000<br>*Test:* 1,000 | (19) |
| **taxonomy_species** | Taxonomic classification at the species level.<br>*Labels:* 618 Species | No | *Train:* 265,276<br>*Val:* 10,000<br>*Test:* 10,012 | (51) |
| **number-of-folds** | Protein fold classification (SCOP 1.75).<br>*Labels:* 1,195 Folds | No | *Train:* 12,312<br>*Val:* 736<br>*Test:* 3,244 | (6) |
| **MB** | Protein-metal interaction prediction.<br>*Labels:* 2 (Binder vs. Non-binder) | No | *Train:* 5,068<br>*Val:* 662<br>*Test:* 665 | (71) |
| **DeepLoc-2** | Binary protein localization.<br>*Labels:* 2 (Membrane-bound vs. Soluble) | No | *Train:* 5,473<br>*Val:* 1,335<br>*Test:* 1,729 | (71) |
| **solubility** | Protein solubility prediction.<br>*Labels:* 2 (Soluble vs. Insoluble) | No | *Train:* 55,536<br>*Val:* 6,942<br>*Test:* 6,942 | (6) |
| **taxonomy_different** | Differential phylogeny prediction.<br>*Labels:* 2 (Same Species vs. Different) | Yes | *Train:* 100,000<br>*Val:* 1,000<br>*Test:* 1,000 | (51) |
| **temp-stability** | Structural stability at 65°C.<br>*Labels:* 2 (Stable vs. Unstable) | No | *Train:* 283,057<br>*Val:* 62,973<br>*Test:* 73,205 | (6) |
| **mat-production** | Suitability for material production.<br>*Labels:* 2 (Suitable vs. Fail) | No | *Train:* 18,541<br>*Val:* 4,786<br>*Test:* 4,790 | (6) |
| **cloning-clf** | Success prediction for the cloning stage.<br>*Labels:* 2 (Success vs. Failure) | No | *Train:* 18,577<br>*Val:* 4,789<br>*Test:* 4,787 | (6) |
| **ecoli_expression** | *E. coli* codon-level expression prediction.<br>*Labels:* 2 (High expression vs. Low) | No | *Train:* 3,424<br>*Val:* 857<br>*Test:* 2,529 | (76) |
| **soluprot** | SoluProt solubility prediction in *E. coli*.<br>*Labels:* 2 (Soluble vs. Insoluble) | No | *Train:* 11,436<br>*Val:* -<br>*Test:* 3,100 | (77) |
| **plm-interact** | Human PPI with cross-species evaluation.<br>*Labels:* 2 (Interaction vs. None) | Yes | *Train:* 421,792<br>*Val:* 52,725<br>*Test:* 242,000 | (78) |
| **gold-ppi** | Leakage-free “gold standard” PPI dataset.<br>*Labels:* 2 (Interaction vs. None) | Yes | *Train:* 163,192<br>*Val:* 59,260<br>*Test:* 52,048 | (52, 53) |
| **KSMoFinder** | Kinase-substrate motif classification (from original study).<br>*Labels:* 2 (Phosphorylates or not) | Yes | *Train:* 72,021<br>*Val:* -<br>*Test:* 6,855 | (79) |
| **KSMoFinder-clustered** | Kinase-substrate motif classification (MMseqs2, 50% similarity at 80% coverage, C3 split).<br>*Labels:* 2 (Phosphorylates or not) | Yes | *Train:* 49,206<br>*Val:* 1,012<br>*Test:* 1,014 | (79) |
| **fitness-prediction** | Protein fitness of GB1 mutants at four specific locations.<br>*Target:* Fitness score | No | *Train:* 6,289<br>*Val:* 699<br>*Test:* 1,745 | (6) |
| **stability-pred** | Protease concentration for structural maintenance.<br>*Target:* Concentration | No | *Train:* 53,614<br>*Val:* 2,512<br>*Test:* 12,851 | (6) |
| **enzyme-*k*<sub>*cat*</sub>** | Enzyme catalytic rate (*k*<sub>cat</sub>).<br>*Target:* Rate value | No | *Train:* 6,837<br>*Val:* 498<br>*Test:* 469 | (6) |
| **optimal-temperature** | Optimal temperature prediction.<br>*Target:* Temperature | No | *Train:* 1,516<br>*Val:* 190<br>*Test:* 190 | (6) |
| **optimal-ph** | Optimal pH prediction.<br>*Target:* pH value | No | *Train:* 7,124<br>*Val:* 760<br>*Test:* 1,971 | (6, 42) |
| **million_full** | Activity of O-methyltransferase variants.<br>*Target:* Activity level | No | *Train:* 93,244<br>*Val:* 11,656<br>*Test:* 11,656 | (80) |
| **PPA-ppi** | Protein-protein affinity prediction. Aggregated from three sources: Bindwell APPT, the Haddock binding affinity benchmark (filtered to two-chain PDB entries), and the Affinity Benchmark v5.5. Clustered at 90% identity (CD-HIT, *n*=3). Sequences were extracted from PDB structures.<br>*Target:* *pK*<sub>*d*</sub> affinity score | Yes | *Train:* 10,639<br>*Val:* 200<br>*Test:* 200 | (81–83) |

**Table S7.**
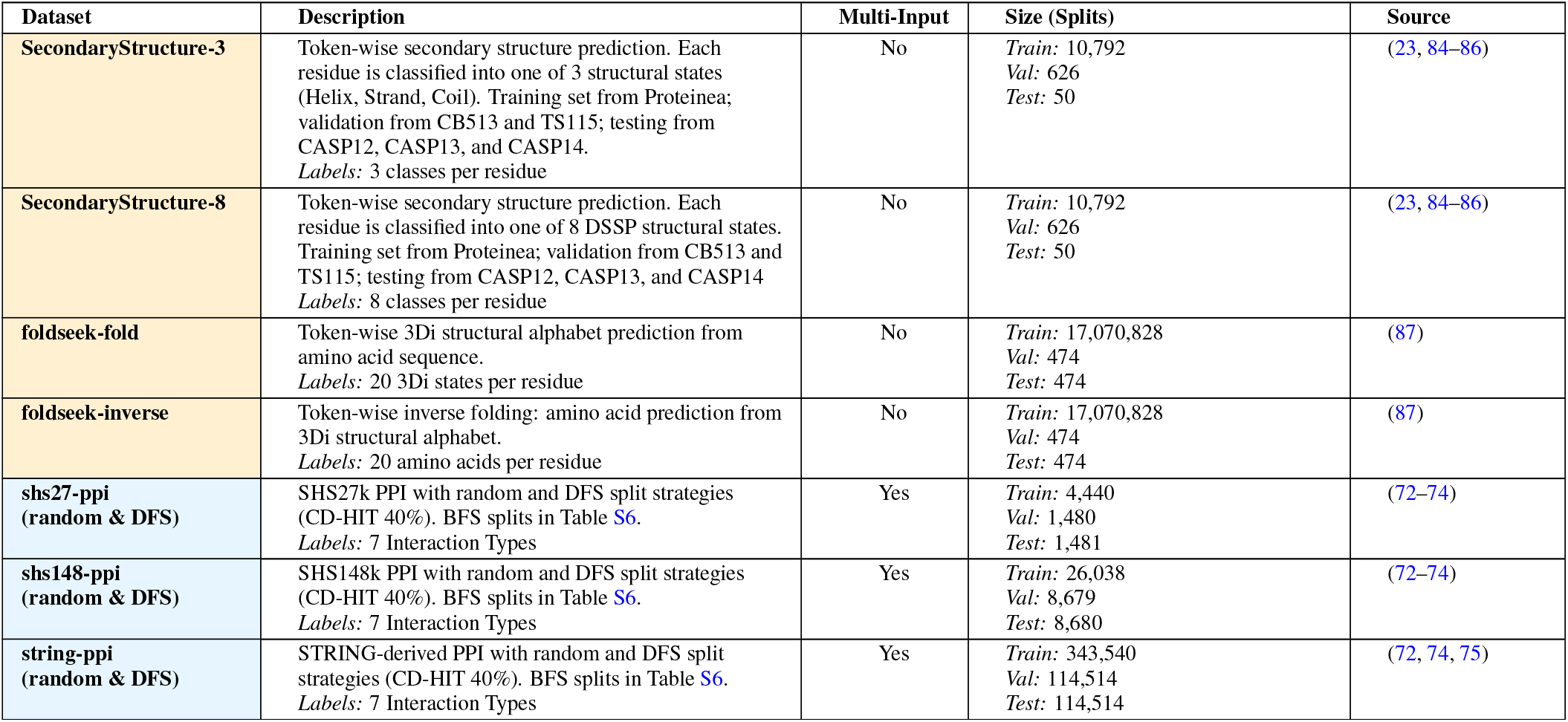
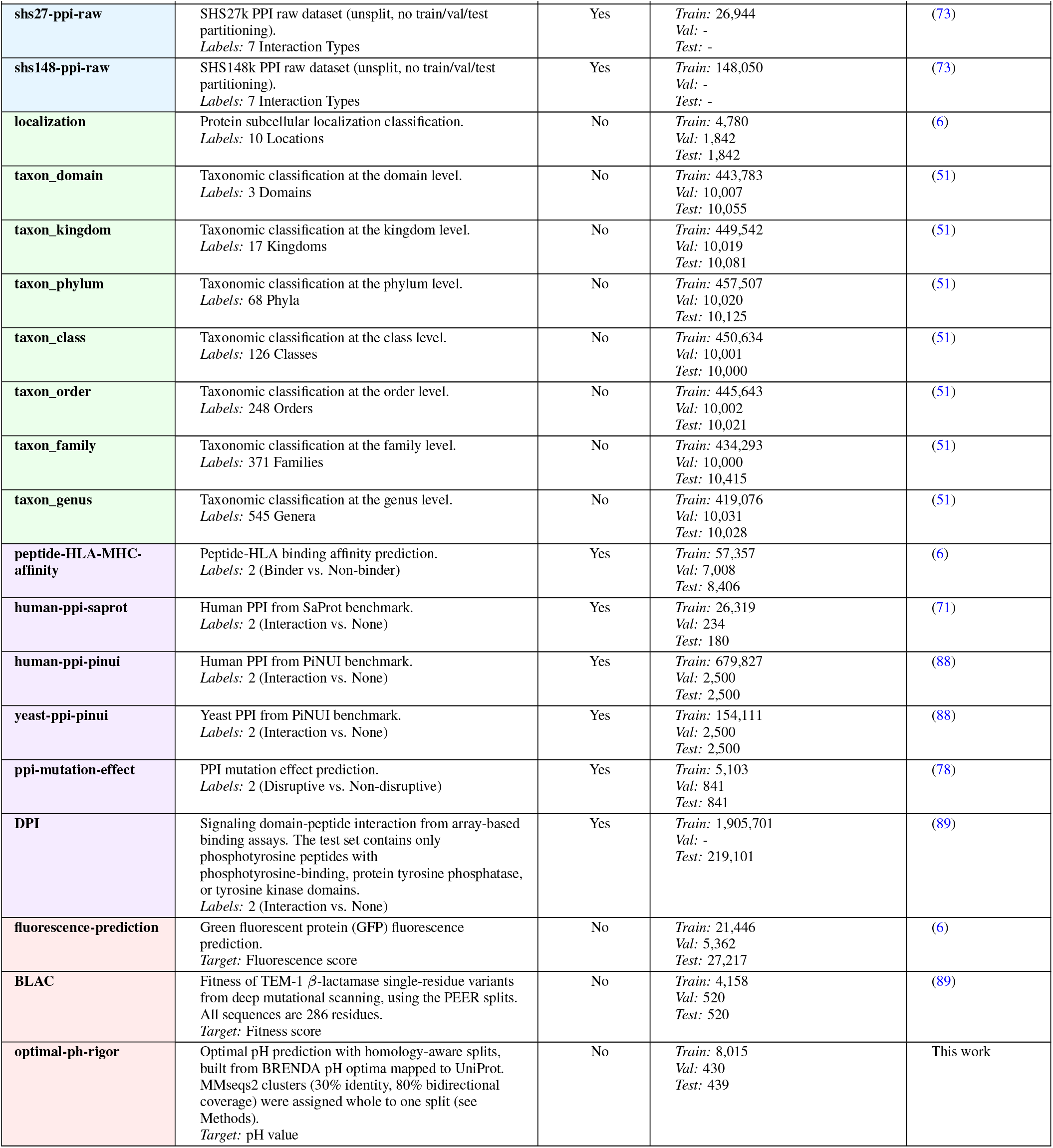
Additional Protify datasets not included in the vector benchmark (Table S6). **Legend:** Token-wise tasks in ; Multi-Label in ; Multi-Class in ; Binary Classification in ; and Regression in .

**Table S8.**
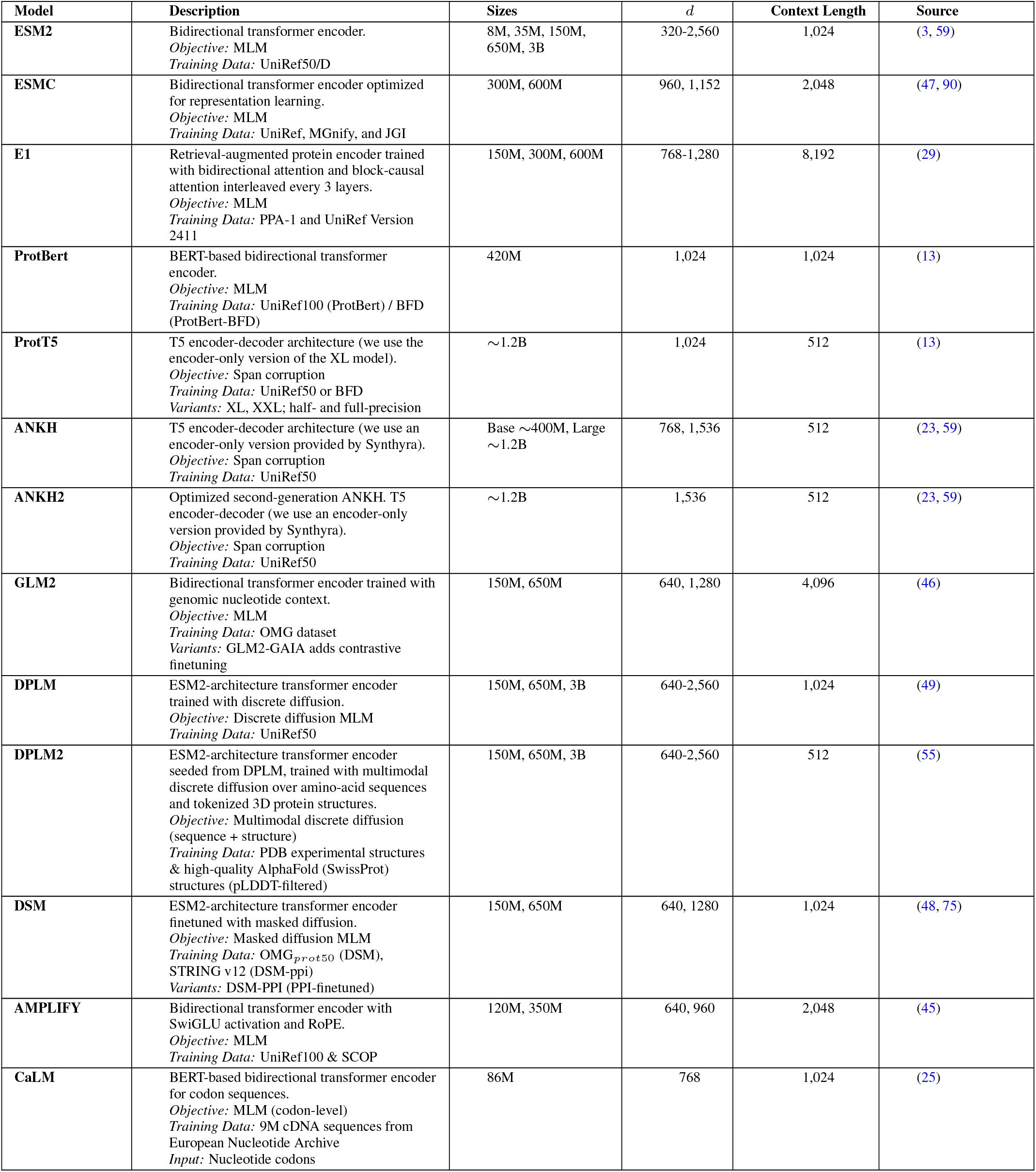
Pretrained models supported by Protify. For each model family, we list available parameter sizes, embedding dimension (*d*), and maximum context length in tokens. Protify also includes three baseline controls not listed here: **Random** (random normal vectors per residue), **OneHot** (one-hot amino acid encoding), and **Random-Transformer** (randomly initialized ESM235M); see Methods for details.

| **Model** | **Description** | **Sizes** | *d* | **Context Length** | **Source** |
|---|---|---|---|---|---|
| **ESM2** | Bidirectional transformer encoder.<br>*Objective:* MLM<br>*Training Data:* UniRef50/D | 8M, 35M, 150M, 650M, 3B | 320-2,560 | 1,024 | (3, 59) |
| **ESMC** | Bidirectional transformer encoder optimized for representation learning.<br>*Objective:* MLM<br>*Training Data:* UniRef, MGnify, and JGI | 300M, 600M | 960, 1,152 | 2,048 | (47, 90) |
| **E1** | Retrieval-augmented protein encoder trained with bidirectional attention and block-causal attention interleaved every 3 layers.<br>*Objective:* MLM<br>*Training Data:* PPA-1 and UniRef Version 2411 | 150M, 300M, 600M | 768-1,280 | 8,192 | (29) |
| **ProtBert** | BERT-based bidirectional transformer encoder.<br>*Objective:* MLM<br>*Training Data:* UniRef100 (ProtBert) / BFD (ProtBert-BFD) | 420M | 1,024 | 1,024 | (13) |
| **ProtT5** | T5 encoder-decoder architecture (we use the encoder-only version of the XL model).<br>*Objective:* Span corruption<br>*Training Data:* UniRef50 or BFD<br>*Variants:* XL, XXL; half- and full-precision | ∼1.2B | 1,024 | 512 | (13) |
| **ANKH** | T5 encoder-decoder architecture (we use an encoder-only version provided by Synthyra).<br>*Objective:* Span corruption<br>*Training Data:* UniRef50 | Base ∼400M, Large ∼1.2B | 768, 1,536 | 512 | (23, 59) |
| **ANKH2** | Optimized second-generation ANKH. T5 encoder-decoder (we use an encoder-only version provided by Synthyra).<br>*Objective:* Span corruption<br>*Training Data:* UniRef50 | ∼1.2B | 1,536 | 512 | (23, 59) |
| **GLM2** | Bidirectional transformer encoder trained with genomic nucleotide context.<br>*Objective:* MLM<br>*Training Data:* OMG dataset<br>*Variants:* GLM2-GAIA adds contrastive finetuning | 150M, 650M | 640, 1,280 | 4,096 | (46) |
| **DPLM** | ESM2-architecture transformer encoder trained with discrete diffusion.<br>*Objective:* Discrete diffusion MLM<br>*Training Data:* UniRef50 | 150M, 650M, 3B | 640-2,560 | 1,024 | (49) |
| **DPLM2** | ESM2-architecture transformer encoder seeded from DPLM, trained with multimodal discrete diffusion over amino-acid sequences and tokenized 3D protein structures.<br>*Objective:* Multimodal discrete diffusion (sequence + structure)<br>*Training Data:* PDB experimental structures & high-quality AlphaFold (SwissProt) structures (pLDDT-filtered) | 150M, 650M, 3B | 640-2,560 | 512 | (55) |
| **DSM** | ESM2-architecture transformer encoder finetuned with masked diffusion.<br>*Objective:* Masked diffusion MLM<br>*Training Data:* OMG<sub>*prot50*</sub> (DSM), STRING v12 (DSM-ppi)<br>*Variants:* DSM-PPI (PPI-finetuned) | 150M, 650M | 640, 1280 | 1,024 | (48, 75) |
| **AMPLIFY** | Bidirectional transformer encoder with SwiGLU activation and RoPE.<br>*Objective:* MLM<br>*Training Data:* UniRef100 & SCOP | 120M, 350M | 640, 960 | 2,048 | (45) |
| **CaLM** | BERT-based bidirectional transformer encoder for codon sequences.<br>*Objective:* MLM (codon-level)<br>*Training Data:* 9M cDNA sequences from European Nucleotide Archive<br>*Input:* Nucleotide codons | 86M | 768 | 1,024 | (25) |

### Supplementary Note 8: Transformer probe hyperparameter sweep

Hyperparameters for the pHopt transformer probe were selected by Bayesian optimization through Protify’s Weights & Biases integration over 50 trials, about four hours of GH200 time. Table S9 reports the search space and the configuration selected.

**Table S9.**
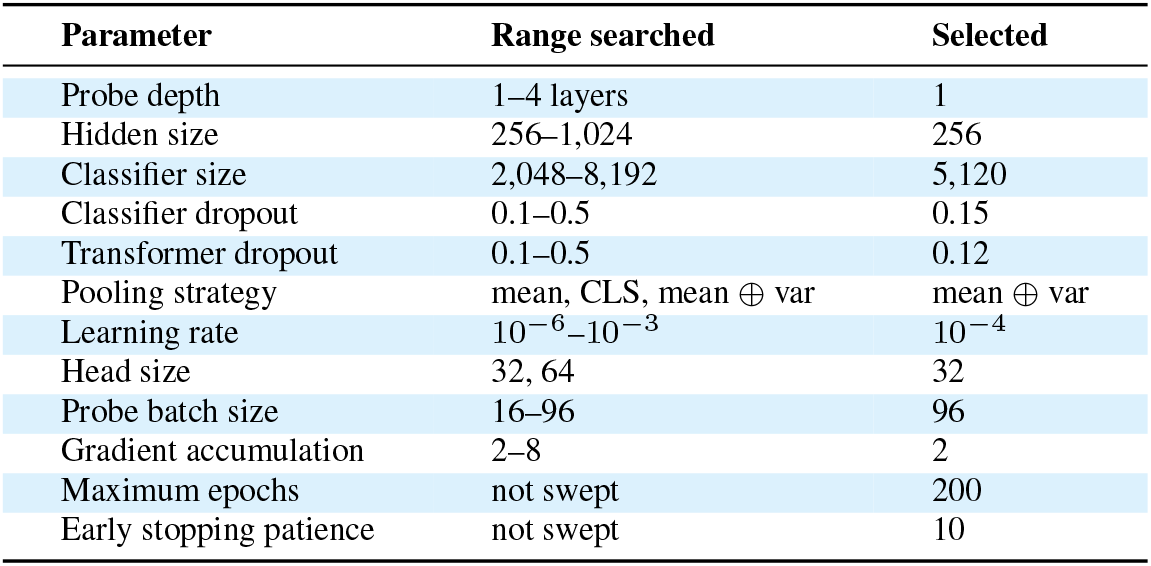
Transformer probe hyperparameter sweep for enzyme optimal pH prediction (DPLM23B per-residue embeddings; 50 trials of Bayesian optimization; objective: validation *R*^2^).

